# Phosphorylation chemistry of the *Bordetella* PlrSR TCS and its contribution to bacterial persistence in the lower respiratory tract

**DOI:** 10.1101/2022.10.06.511181

**Authors:** Sarah A. Barr, Emily N. Kennedy, Liliana S. McKay, Richard M. Johnson, Ryan J. Ohr, Peggy A. Cotter, Robert B. Bourret

## Abstract

2

*Bordetella* species cause lower respiratory tract infections in mammals. *B. pertussis* and *B. bronchiseptica* are the causative agents of whooping cough and kennel cough, respectively. The current acellular vaccine for *B. pertussis* protects against the pertussis toxin but does not prevent transmission or colonization. Cases of *B. pertussis* infections are on the rise even in areas of high vaccination. The PlrSR two-component system, is required for persistence in the mouse lung. A partial *plrS* deletion strain and a *plrS H521Q* strain cannot survive past three days in the lung, suggesting PlrSR works in a phosphorylation dependent mechanism. We characterized the biochemistry of *B. bronchiseptica* PlrSR and found that both proteins function as a canonical two-component system. His521 and Glu522 were essential for PlrS autophosphorylation. Asn525 was essential for phosphatase activity. The PAS domain was critical for both PlrS autophosphorylation and phosphatase activities. PlrS can both phosphotransfer to and exert phosphatase activity towards PlrR. Unexpectedly, PlrR forms a tetramer when unphosphorylated and a dimer upon phosphorylation. Finally, we demonstrated the importance of PlrS phosphatase activity for persistence within the murine lung. By characterizing PlrSR we hope to guide future *in vivo* investigation for development of new vaccines and therapeutics.

**GRAPHICAL ABSTRACT:** 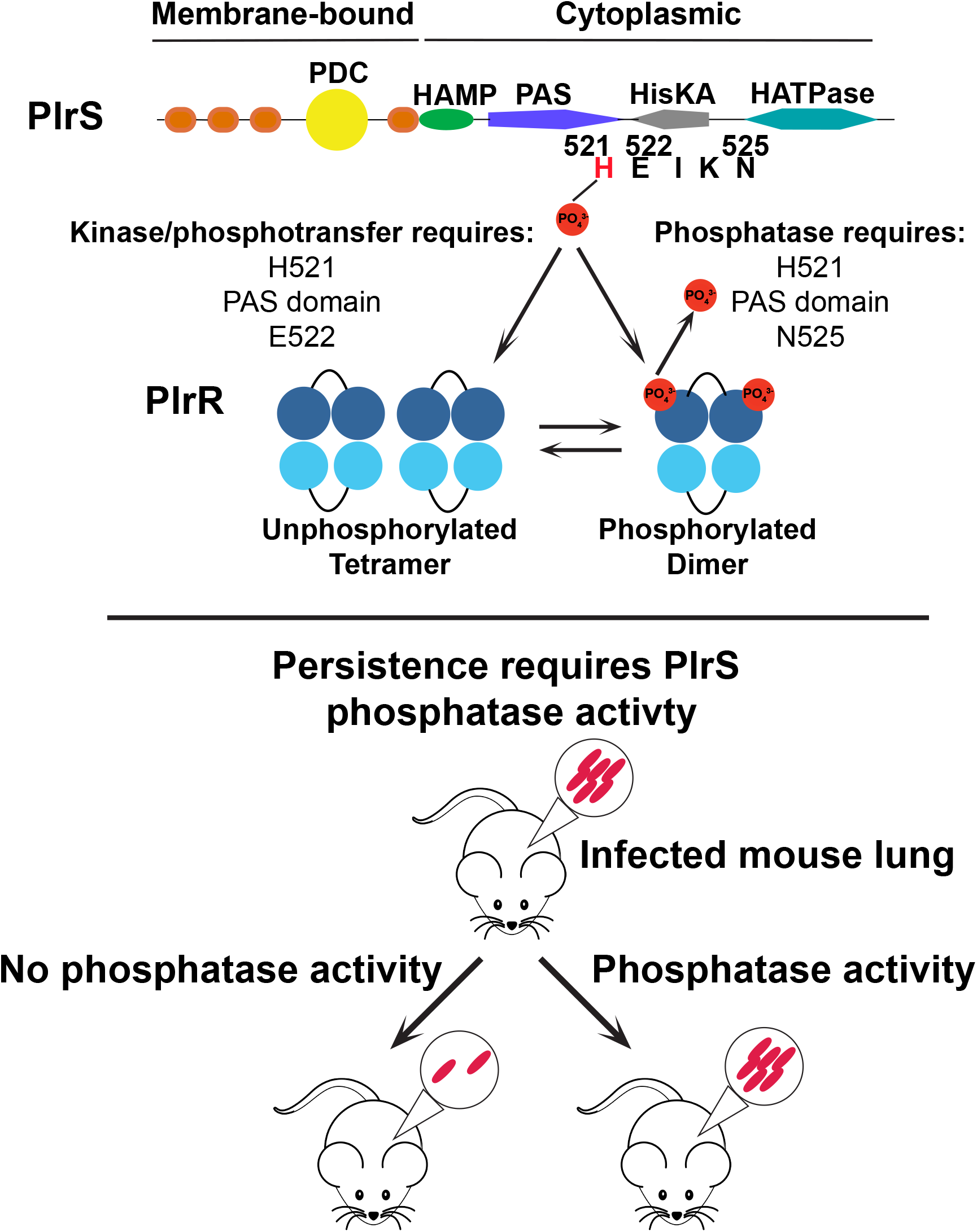

## 4 INTRODUCTION

Bacteria are exposed to a multitude of changes in nutrients, pH, viscosity, temperature etc. Although bacteria residing in communities can be buffered from some harsh changes, survival of individual bacterial cells requires efficient sensing and adaption to ever-changing environments. For example, swimming toward a chemoattractant, upregulation of virulence mechanisms upon host invasion, and secretion of protective capsules to avoid phagocytosis all require dynamic adaptation. A diverse category of signaling proteins called two-component systems (TCSs) allow bacterial cells to sense dynamic changes in their environment and integrate those signals into corresponding outputs on a wide range of time scales (Zschiedrich *et al*., 2016); chemotaxis takes place within seconds (Segall *et al*., 1982), while sporulation takes place over hours (Piggot & Hilbert, 2004). The most common type of bacterial TCS is composed of a membrane bound sensor kinase that senses an external stimulus and encodes a resultant signal via autophosphorylation. The phosphoryl group is transferred to a cytoplasmic partner response regulator whose activity is modulated by phosphorylation. The response regulator carries out a corresponding biological response on an appropriate time scale (Buschiazzo & Trajtenberg, 2019). Most bacterial response regulators control gene expression, however there are many other regulatory targets such as second messengers (*e.g*., cyclic-di-GMP) or the flagellar motor (Galperin, 2010).

The mechanisms by which TCSs acquire, transfer, and lose phosphoryl groups are preserved across systems via conserved residues that carry out similar chemistry in different contexts, although, many characteristics distinguish TCSs from one another and help define specialized biological functions. For example, the phosphorylation chemistry of the conserved histidine in sensor kinases is influenced by adjacent amino acids (Huynh & Stewart, 2011; Multamaki *et al*., 2021); the E/D residue at the H+1 position is typically critical for supporting autophosphorylation (Atkinson & Ninfa, 1993; Willett & Kirby, 2012), whereas the N/T residue at the H+4 position is typically critical for coordinating the attacking water molecule to support dephosphorylation of the phosphorylated response regulator (*i.e*., phosphatase activity) (Atkinson & Ninfa, 1993; Dutta *et al*., 2000; Hsing *et al*., 1998; Pazy *et al*., 2010; Silversmith, 2010; Willett & Kirby, 2012). The balance between autophosphorylation, phosphotransfer, and phosphatase activity toward the response regulator defines how frequently the system is active or inactive. Furthermore, the speed at which the system can acquire and transfer or remove phosphoryl groups defines the response time to a stimulus as well as duration of the response.

Bacteria of the genus *Bordetella* cause respiratory tract infections in mammals. *Bordetella bronchiseptica*, which has a broad host range, is a close ancestor of the obligate human pathogen *Bordetella pertussis*, the causal agent of whooping cough. The two species are highly similar genetically and produce a nearly identical set of virulence factors, many of which are functionally interchangeable (Martinez de Tejada *et al*., 1996). Currently used acellular *B. pertussis* vaccines induce antibody responses that prevent disease symptoms by neutralizing virulence factors, but do not prevent bacterial colonization or transmission (Warfel *et al*., 2014), resulting in spread to unvaccinated individuals and the formation of a *B. pertussis* reservoir that can mutate to escape vaccines. Further, acellular pertussis vaccines do not support long-term immunity and require an increasing number of boosters (Esposito *et al*., 2019). Consequently, *B. pertussis* cases are trending higher even in geographical areas with a high vaccination rate (Decker & Edwards, 2021; Fry *et al*., 2021; Yeung *et al*., 2017)

The BvgAS (*<u>B</u>ordetella* <u>v</u>irulence <u>g</u>ene) TCS is considered the master virulence regulator in bordetellae and controls the expression of hundreds of genes, including many of those encoding known protein virulence factors (Belcher *et al*., 2020; Cotter & Miller, 1997; Hot *et al*., 2003; Lacey, 1960; Moon *et al*., 2017; Weiss & Falkow, 1984). BvgAS activity is necessary for respiratory infection; BvgAS null mutants are avirulent, mutants with constitutively active BvgAS systems are indistinguishable from wild-type bacteria *in vivo*, and mutants that fail to repress BvgAS-repressed genes are defective for infection (Akerley *et al*., 1995; Cotter & Miller, 1994; Martinez de Tejada *et al*., 1998; Merkel *et al*., 1998). However, another TCS, named PlrSR (<u>p</u>ersistence in the <u>l</u>ower <u>r</u>espiratory tract, <u>s</u>ensor kinase and <u>r</u>esponse regulator), is critical for both bacterial persistence and BvgAS activity in the lower respiratory tract (LRT) (Bone *et al*., 2017; Kaut *et al*., 2011; Sobran & Cotter, 2019).

PlrS and PlrR share substantial predicted amino acid similarity with NtrY and NtrX from betaproteobacteria. Like its *ntrX* homologs (Bonato *et al*., 2016; Gregor *et al*., 2007; Ishida *et al*., 2002), *plrR* is essential in *B. bronchiseptica* and *B. pertussis. B. bronchiseptica* strains with plasmid insertions in *plrS*, small deletions at the 5’ end of *plrS*, or encoding PlrS proteins in which the predicted site of phosphorylation is replaced with glutamine (H521Q) are cleared from the trachea and lungs of mice within a few days post-inoculation, indicating that PlrS phosphorylation, and hence presumably PlrR phosphorylation, is required for bacterial survival in the LRT (Bone *et al*., 2017; Kaut *et al*., 2011). *B. pertussis* with a partial deletion in *plrS* is also defective for persistence in the LRT in mice (Bone *et al*., 2017). All phenotypes identified for *plrS* mutants *in vitro* so far are also BvgAS-regulated phenotypes, and we have shown that BvgAS activity in the LRT requires *plrS*. However, *plrS* mutants are also unable to persist in the LRT when BvgAS is rendered constitutively active by an amino acid substitution in the BvgS linker (Bone *et al*., 2017), indicating that PlrSR must either activate or repress one or more BvgAS-independent genes for the bacteria to survive in the LRT. Gene products that are both PlrSR-dependent and BvgAS-independent genes could serve as therapeutic targets or new vaccine components and hence their identification is a major goal. Identifying PlrSR-regulated genes is currently hampered, however, by the facts that: *plrR* is essential; the active conformation (phosphorylated or unphosphorylated or both) of PlrR is unknown; conditions in which PlrS is active or inactive *in vitro* are unknown; mutants in which PlrS or PlrR are definitively inactive or constitutively active are unknown; and PlrSR-dependent, BvgAS-independent phenotypes are unknown.

Here, we utilized a biochemical approach to gain access to the PlrSR TCS. We determined the multimeric state and phosphoryl group stability of PlrR, as well as the impact of the PlrR DNA binding domain on phosphotransfer reactions. We determined the contributions of the H+1 and H+4 residues as well as PlrS regulatory domains to (i) PlrS autophosphorylation kinetics, (ii) the balance between PlrS kinase and phosphatase activity, and (iii) phosphotransfer and phosphatase kinetics of PlrS toward PlrR. Based on the results of these *in vitro* experiments, we constructed a *B. bronchiseptica* strain producing a PlrS protein defective for phosphatase activity and found that both kinase and phosphatase functions of PlrS are critical for persistence in the LRT.

## 5 RESULTS

### *Overview of* B. bronchiseptica *PlrS structure*

PlrS is a putative membrane-bound, sensor histidine kinase (Figure 1). PlrS contains four predicted transmembrane domains, a periplasmic PhoQ-DcuS-CitA (PDC) domain, and cytoplasmic HAMP, PAS, HisKA, and HATPase_c domains. To avoid issues of solubility or protein folding, only cytoplasmic portions of PlrS were used in biochemical experiments. HAMP domains use conformational changes to transmit input signals to the HisKA (dimerization and histidine phosphorylation) and HATPase_c (catalytic and ATP binding) domains to regulate autophosphorylation (Parkinson, 2010). PAS domains are sensory domains that can work in tandem with, or independently of, other sensory domain(s) of the sensor kinase. PAS domains are structurally conserved but diverse both in the specific ligands they bind and mechanisms for regulation of phosphorylation chemistry within a sensor kinase (Henry & Crosson, 2011; Stuffle *et al*., 2021).

**Figure 1.**
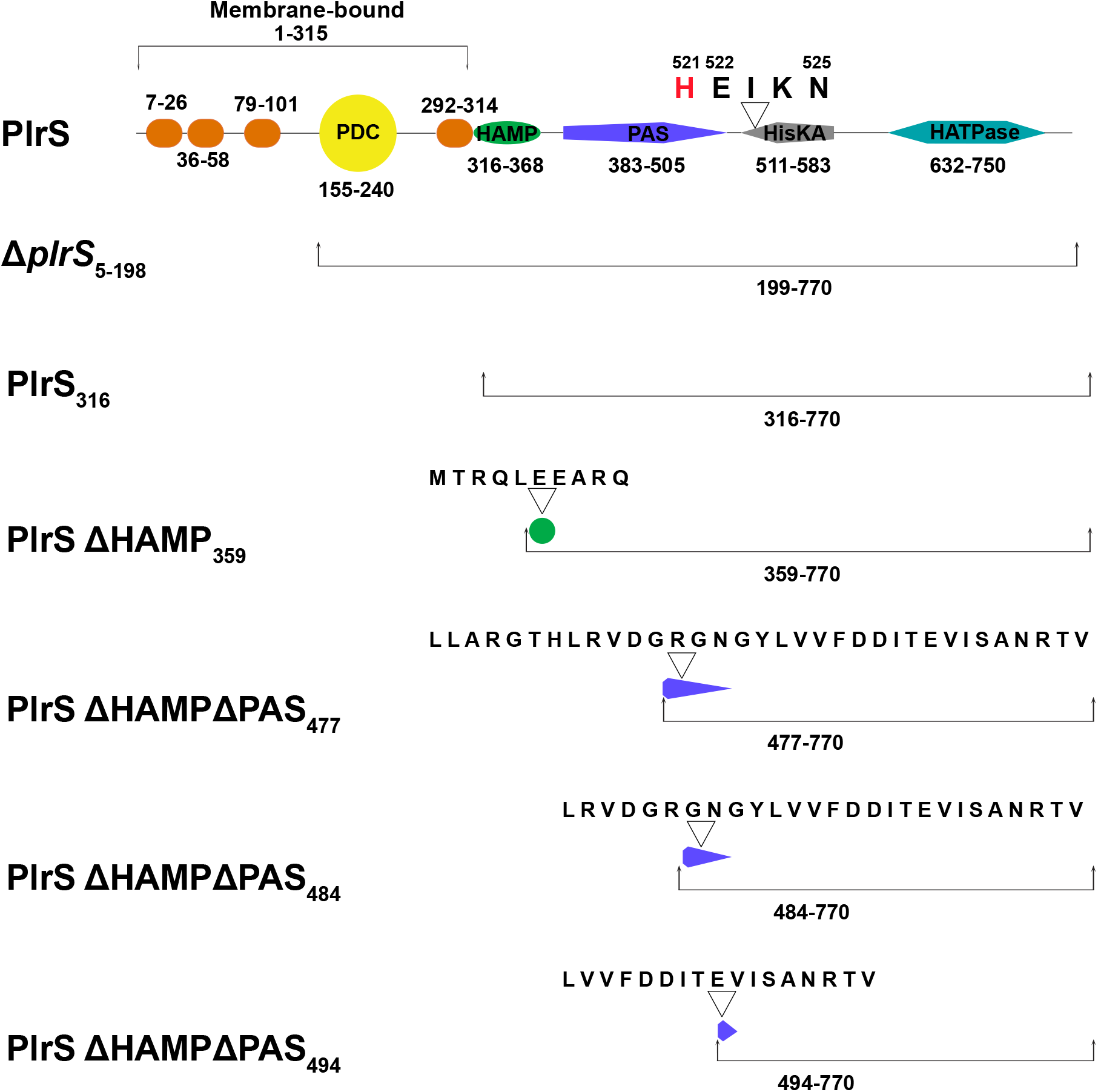
PlrS proteins used in this study. Full length PlrS (top) from *B*. *bronchiseptica* is predicted to be a membrane bound sensor kinase with a periplasmic PDC sensory domain and cytoplasmic HAMP, PAS, HisKA, and HATPase_c domains. Domains and their amino acid boundaries identified by the SMART database (Letunic *et al*., 2021) are labeled. The proteins used in this study are organized by size and are annotated by the N-terminal residue relative to full-length PlrS. Some proteins contained amino acid substitutions at positions 521, 522, or 525. For each truncated mutant we retained a small number of amino acids N-terminal to the predicted junction. The remaining amino acid sequences are indicated above the representative truncated domain cartoon.

The HisKA domain contains the conserved His (position 521) predicted to be phosphorylated in response to stimulus recognition. The H+1 position at 522 and the H+4 position at 525 are Glu and Asn, respectively. We used proteins with single amino acid substitutions at these positions to probe the phosphorylation chemistry mechanisms of PlrS. The mutants are named by their N-terminal residue in subscript (*i.e*., PlrS_316_) followed by the amino acid substitution (*i.e*., N525A). To probe the contribution of the HAMP and PAS domains to phosphorylation chemistry of PlrS, we used mutant proteins lacking all sequences N-terminal to the respective domains and the domain itself (Figure 1). For each truncated protein, a small number of amino acids N-terminal to the predicted domain junction were retained to allow for efficient structural folding. We used a nested set of PlrS ΔHAMPΔPAS proteins in an attempt to avoid disruption of the adjacent HisKA domain. The protein encoded by the Δ*plrS*_5-198_ allele was not purified for biochemical studies and it refers to the previously characterized mutant (Kaut *et al*., 2011) used for *in vivo Bordetella* experiments.

### *PlrS functions as a kinase* in vitro

To determine if PlrS functions as a histidine kinase, we conducted autophosphorylation experiments in which 10 μM of each PlrS protein was incubated with 100 μM [γ-^32^P]ATP for 15 minutes at room temperature. PlrS_316_ autophosphorylated, as expected for a sensor histidine kinase (Figure 2). PlrS_316_ H521Q did not incorporate radiolabel, consistent with His521 serving as the only site of phosphorylation. PlrS_316_ E522A incorporated much less label than PlrS_316_, suggesting the H+1 Glu is critical for efficient autophosphorylation (Willett & Kirby, 2012). PlrS_316_ N525A incorporated label at slightly reduced levels compared with PlrS_316_. The two PlrS ΔHAMP_359_ proteins incorporated radiolabel, whereas removal of the HAMP and PAS domains resulted in undetectable autophosphorylation activity within the 15-minute experiment. The failure to observe autophosphorylation of PlrS proteins lacking the HAMP and PAS domains was consistent across two separate protein purifications of both PlrS ΔHAMPΔPAS_494_ and PlrS ΔHAMPΔPAS_494_ N525A proteins. Observation of autophosphorylation for PlrS ΔHAMP but not PlrS ΔHAMPΔPAS proteins indicate that the PAS domain is critical for PlrS autophosphorylation within the context of this experimental design.

**Figure 2.**
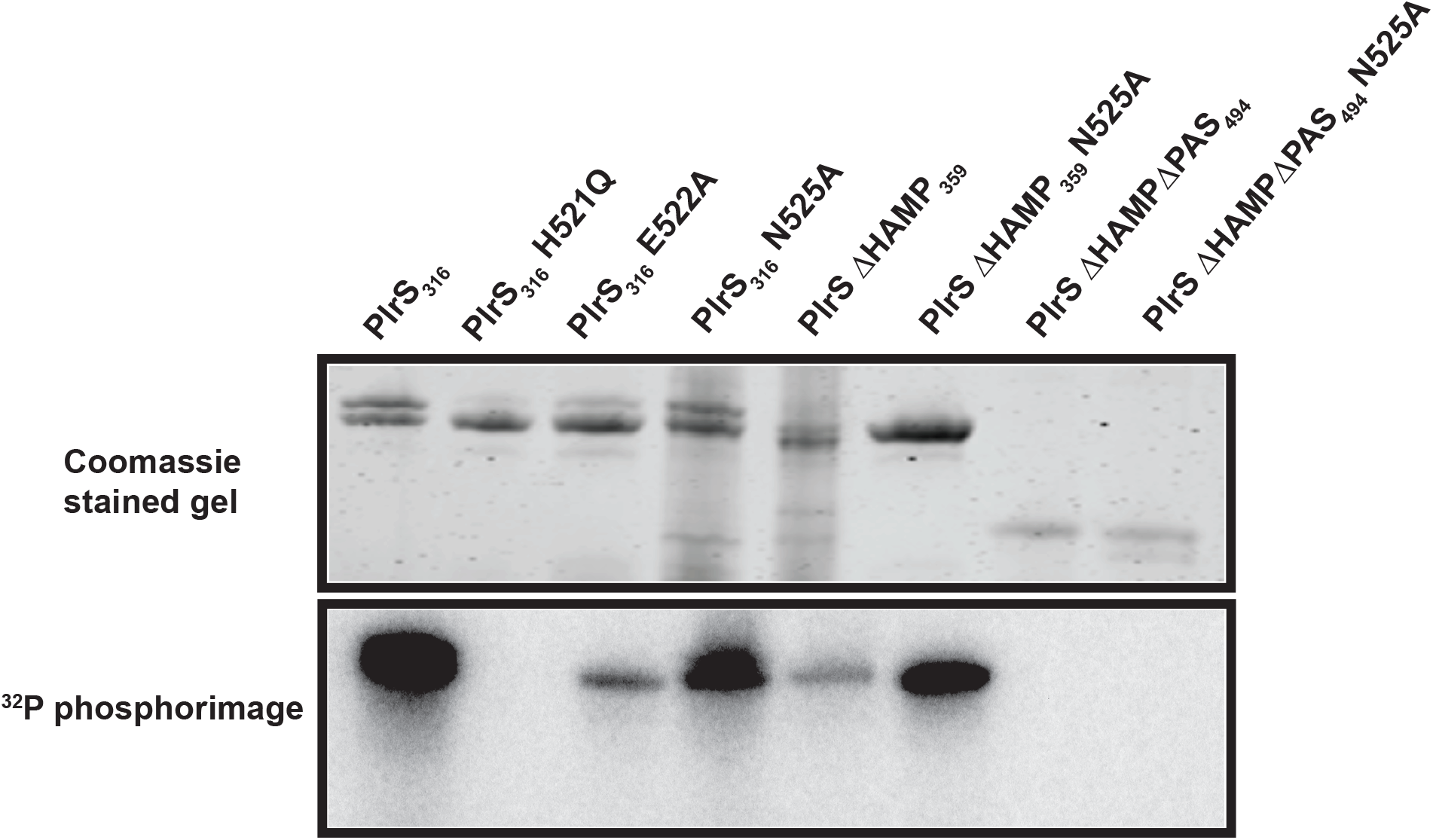
PlrS proteins autophosphorylated to different levels depending on which domains were present. PlrS proteins were incubated with [γ-^32^P]ATP. After 15 minutes the reaction was quenched with stop buffer. Top: PlrS proteins separated using SDS-PAGE followed by staining with Coomassie brilliant blue. Bottom: Phosphorimage of PlrS proteins after a 15-minute incubation. Experiment was performed in triplicate; one representative experiment is shown.

### The PAS domain is critical for PlrS autophosphorylation

To better determine the contribution of key PlrS residues and domains (Figure 1) to the rate of autophosphorylation, we conducted autophosphorylation time courses with wild-type and mutant PlrS proteins (Figure 3). Autophosphorylation reactions were initiated by adding 100 μM [γ-^32^P]ATP to 10 μM PlrS. Aliquots were removed and reactions halted by mixing with Laemmli sample buffer at various time points. PlrS_316_ exhibited such fast autophosphorylation at room temperature that we could not manually measure the initial phase of the reaction, so experiments were conducted at 0 °C.

**Figure 3.**
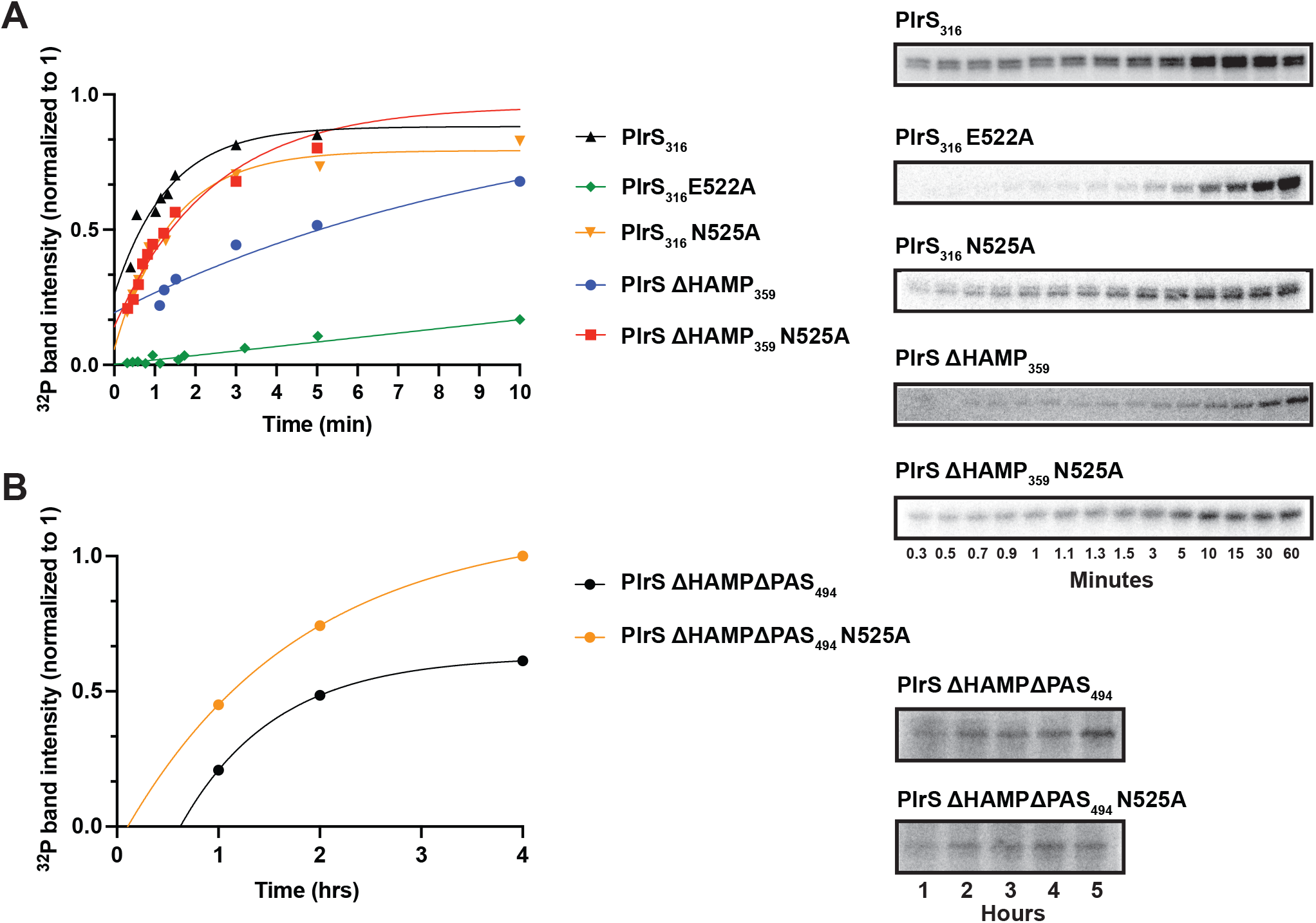
PlrS autophosphorylation kinetics. Representative phosphorimages (right) and data plots (left) of PlrS autophosphorylation experiments. A. To measure rate constants, autophosphorylation of most PlrS proteins were conducted at 0 °C over 1 hour. B. Autophosphorylation experiments for PlrS ΔHAMPΔPAS_494_ and PlrS ΔHAMPΔPAS_494_ N525A proteins were conducted at room temperature over 5 hours. For all experiments, data were normalized to the maximal value and fit to an exponential decay equation. Experiments were conducted in triplicate.

The PlrS_316_ autophosphorylation rate constant was 0.62 ± 0.4 min^−1^ (n=7) (Table 1). The PlrS_316_ N525A rate constant was 0.94 ± 0.09 min^−1^ (n = 3), corroborating the steady state data that PlrS_316_ N525A has relatively the same autophosphorylation efficiency as PlrS_316_ (Figure 2). The high variation in rate constant for PlrS_316_ might be due to slow stochastic switching between kinase and phosphatase conformations at 0°C. PlrS_316_ E522A showed a dramatic rate constant deficit, again corroborating the steady state data and confirming a critical role for the H+1 position in autophosphorylation. PlrS ΔHAMP_359_ exhibited an intermediate rate constant, suggesting that conformations facilitated by the HAMP domain are important for efficient autophosphorylation. Sensor kinases switch between conformations that favor autophosphorylation or phosphatase activity (Buschiazzo & Trajtenberg, 2019). In the presence of the HAMP domain, the N525A substitution had little effect on PlrS autophosphorylation. However, in the absence of the HAMP domain, proteins bearing Asn525 or Ala525 exhibited different autophosphorylation kinetics (Table 1), potentially suggesting Asn525 stabilized phosphatase conformations and led to slower autophosphorylation.

**Table 1.**
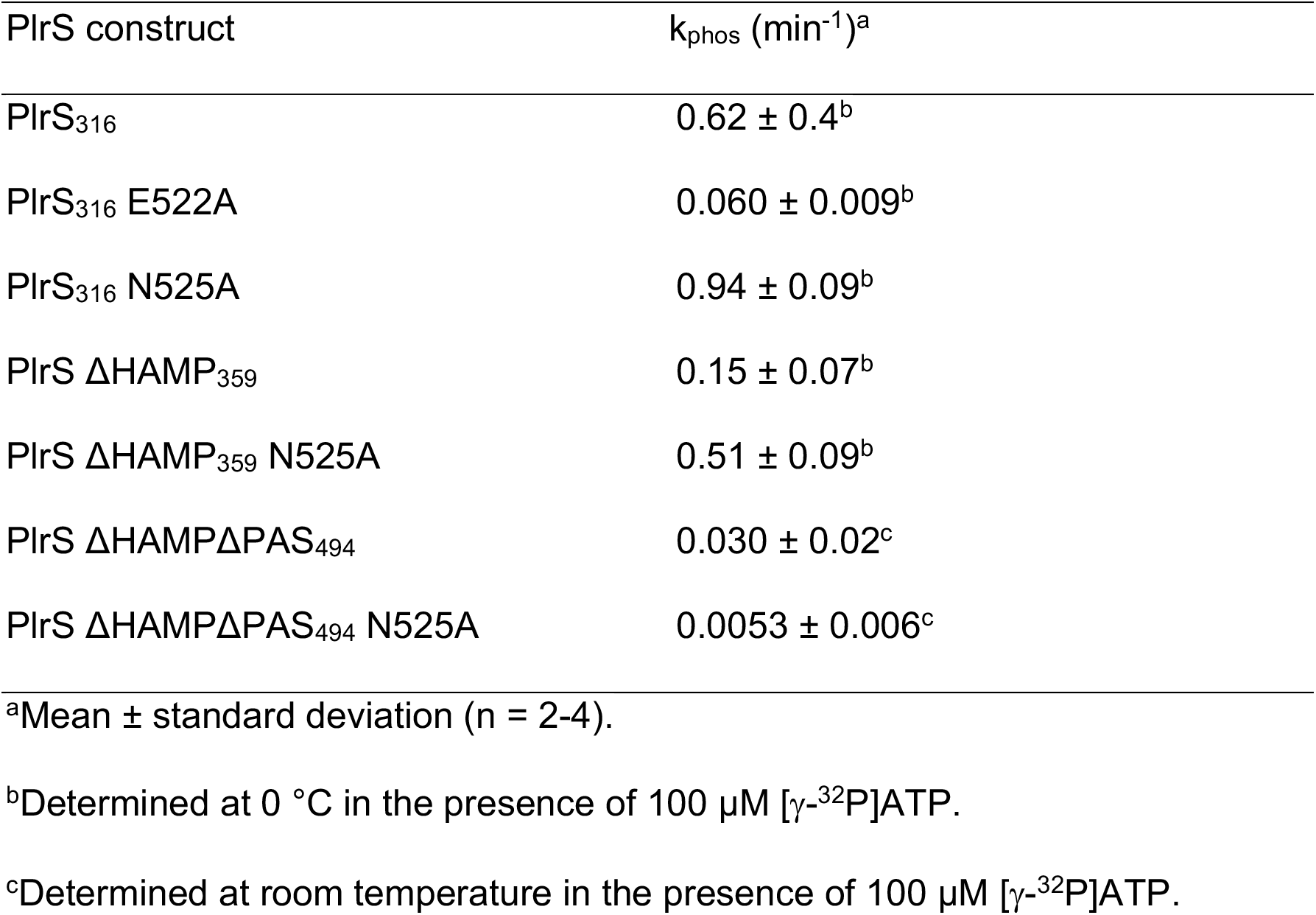
Autophosphorylation rate constants of PlrS proteins.

PlrS ΔHAMPΔPAS_494_ and PlrS ΔHAMPΔPAS_494_ N525A did not exhibit detectable autophosphorylation within one hour at room temperature (data not shown). However, PlrS ΔHAMPΔPAS_494_ and PlrS ΔHAMPΔPAS_494_ N525A did show autophosphorylation over the course of 5 hours with rate constants of 1.8 ± 1 and 0.32 ± 0.4 hr ^−1^ respectively (Figure 3B, Table 1). The observation of slow autophosphorylation suggests that the PAS domain is critical for efficient autophosphorylation and rules out an alternative explanation that the lack of observable autophosphorylation in initial experiments was due to completely nonfunctional proteins. Although the autophosphorylation rate constants of PlrS_316_ and PlrS_316_ N525A suggest Asn525 stabilizes the phosphatase conformation, the opposite pattern was observed with PlrSΔHAMPΔPAS_494_ versus PlrSΔHAMPΔPAS_494_ N525A, perhaps reflecting the different temperatures at which the experiments were conducted and/or the effects of the HAMP and PAS domains on conformational stability. Furthermore, the observed rate constants were highly variable, likely due to a low signal to noise ratio. To assess if the slow rate constants were artifacts of the specific truncation at amino acid 494 chosen for the predicted junction of the HisKA and PAS domains, we also assayed the autophosphorylation rates of two longer proteins (PlrS ΔHAMPΔPAS_477_, PlrS ΔHAMPΔPAS_484_, Figure 1). Both proteins exhibited slow autophosphorylation rate constants similar to PlrS ΔHAMPΔPAS_494_ (Figure S1), increasing confidence in the interpretation that the PAS domain is critical for PlrS autophosphorylation.

### Unphosphorylated PlrR is a multimer that dimerizes upon phosphorylation

To determine the oligomeric state of PlrR, purified protein was subjected to size exclusion chromatography. The calculated molecular mass of monomeric PlrR is 25 kDa. However, PlrR eluted from a calibrated analytical size exclusion column as a broad peak ranging in size from approximately 50 kDa to 90 kDa (data not shown). These data suggest that a heterogeneous population of PlrR multimers were present in our preparation. Additionally, native PlrR did not flow through a filter with a molecular weight cutoff of 100 kDa. To determine the true oligomeric state of purified PlrR, we performed mass photometry. Mass photometry determines the size of a protein based on light scattered by proteins at an interface (Young *et al*., 2018). When PlrR was in solution at room temperature, mass photometry data indicated that PlrR existed as a mixed population of apparent dimer and tetramer species, indicated by the two histogram peaks near 78 and 125 kDa (Figure 4). When PlrR was incubated with the phosphomimic BeF_3_^−^ (Yan *et al*., 1999) (formed *in situ* by combining BeSO_4_ and NaF), the PlrR population existed exclusively as dimers indicated by the single histogram peak at 74/77 kDa (Figure 4). The addition of NaF alone did not affect the distribution of dimer and tetramer, indicating that the observation was BeF_3_^−^ specific. To determine if the change in oligomeric state was activation-dependent or an artifact of the presence of BeF_3_^−^ in the sample, we also assayed a PlrR D52A mutant lacking the predicted Asp site of phosphorylation. PlrR D52A did not shift in population ratio between dimer and tetramer upon addition of BeF_3_^−^ (Figure 4). These data suggest that PlrR exists at an equilibrium between dimer and tetramer when unphosphorylated, but upon BeF_3_^−^ binding (presumably mimicking phosphorylation), the equilibrium shifts in favor of a dimer. The receiver domain alone (PlrR-Rec) (as monomer, dimer, or tetramer) is below the size limit of detection for the mass photometry instrument. However, size exclusion chromatography showed PlrR-Rec as a mixed population between monomer and dimer (data not shown). The difference in oligomeric state between full-length and PlrR-Rec suggests that tetramerization involves the DNA binding domain.

**Figure 4.**
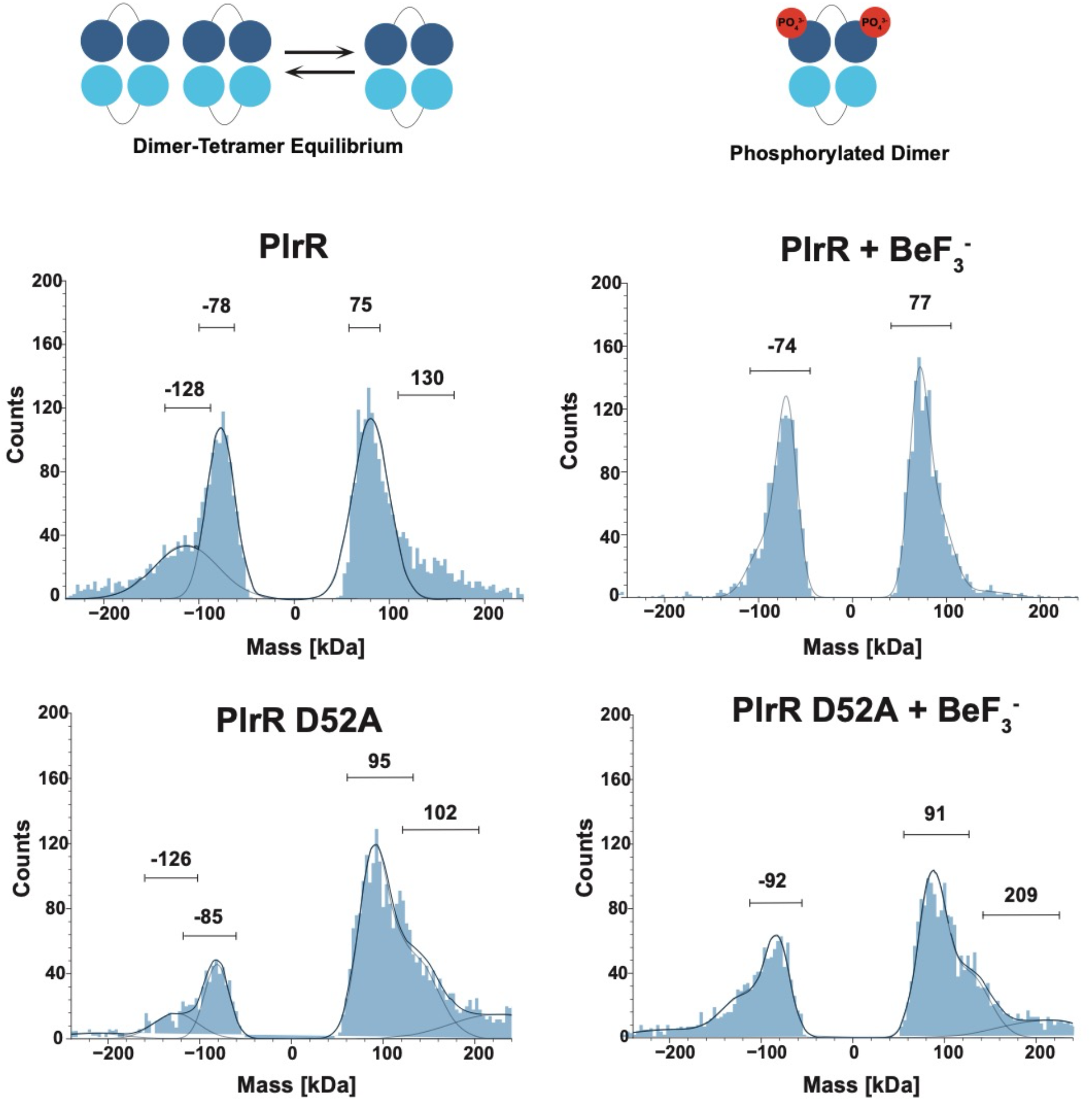
Mass photometry of PlrR. Histogram plots of mass for each impact of a protein particle. Positive mass is a metric of impact while negative mass is a metric of detachment. Plots are of PlrR or PlrR D52A with or without the phosphoryl mimic, BeF_3_^−^.

### Autophosphorylation of PlrR with small molecule phosphodonors was not detected

Many response regulators can catalyze their own phosphorylation using small molecule phosphodonors (Lukat *et al*., 1992; Silversmith *et al*., 1997; Vazquez-Ciros *et al*., 2020; Wolfe, 2010). Changes in fluorescence intensity of a Trp residue upon phosphorylation at a neighboring residue was used to demonstrate autophosphorylation of the *Brucellus abortis* NtrX receiver domain with acetyl phosphate (Fernandez *et al*., 2015). PlrR contains a Trp residue analogous to the NtrX Trp residue located two residues C-terminal to the predicted Asp52 phosphorylation site. However, we were unable to detect autophosphorylation of PlrR-Rec with acetyl phosphate, phosphoramidate, or monophosphoimidizole using Trp fluorescence (data not shown). Furthermore, autophosphorylation of PlrR and PlrR-Rec using [^32^P]acetyl phosphate was also not detectable (data not shown). Control reactions demonstrated successful autophosphorylation of *Escherichia coli* CheY by both fluorescence and radioactivity.

### PlrS serves as a phosphodonor for PlrR

The adjacent location of *plrS* and *plrR*, presumably in an operon, suggests that PlrS and PlrR form a TCS (Kaut *et al*., 2011). To test the prediction that PlrS and PlrR interact as partner sensor kinase and response regulator, a 10 μM solution of each PlrS protein was allowed to autophosphorylate with 100 μM [γ-^32^P]ATP for 15 minutes, then either full length PlrR or the PlrR-Rec domain was added to a final concentration of 10 μM. Each autophosphorylating PlrS protein could donate phosphoryl groups to both PlrR and PlrR-Rec (Figure 5A). When incubated with PlrS_316_ E522A, there was significantly less radiolabel accumulation on PlrR and PlrR-Rec, compared to incubation with PlrS_316_, presumably due to rate limiting autophosphorylation of PlrS_316_ E522A (Table 1). PlrR-Rec rapidly acquired label from PlrS and then lost all label to dephosphorylation, while full-length PlrR maintained a constant amount of label over one hour (Figure 5B). The differences in accumulation of phosphorylated protein observed between full-length PlrR and PlrR-Rec clearly indicated that the PlrR DNA binding domain, and therefore potentially the unphosphorylated tetramer structure, impaired phosphotransfer from PlrS. Because four reactions (PlrS autophosphorylation, phosphotransfer to PlrR, autodephosphorylation of PlrR, and PlrS phosphatase activity toward PlrR) occur simultaneously, it was not obvious whether PlrR-P was a worse substrate than PlrR-Rec-P for the phosphatase activity of PlrS (Figure 5B). We therefore used a computer simulation as described in Experimental Procedures to successfully model the data for experiments involving PlrR-Rec (data not shown). The software could fit the PlrR-Rec data when only k_+1_ (PlrS autophosphorylation) was set to the experimental value of 0.62 ± 0.4 min^−1^ and the other rate constants were allowed to float (Table 1). However, when we applied the same k_+1_ as with PlrR-Rec and constrained k_+3_ (PlrR autodephosphorylation) and k_+4_ (PlrS phosphatase activity) to the rate constants calculated from the PlrR-Rec analysis to the full-length PlrR, the software could not fit the data while allowing only the phosphotransfer rate constant to float (data not shown). However, the PlrR data could be fit if the rate constants for both the phosphotransfer and phosphatase reactions were allowed to assume slower values compared to the reaction with PlrR-Rec (data not shown). We therefore conclude that both phosphotransfer and phosphatase activities of PlrS were slower toward PlrR than PlrR-Rec.

**Figure 5.**
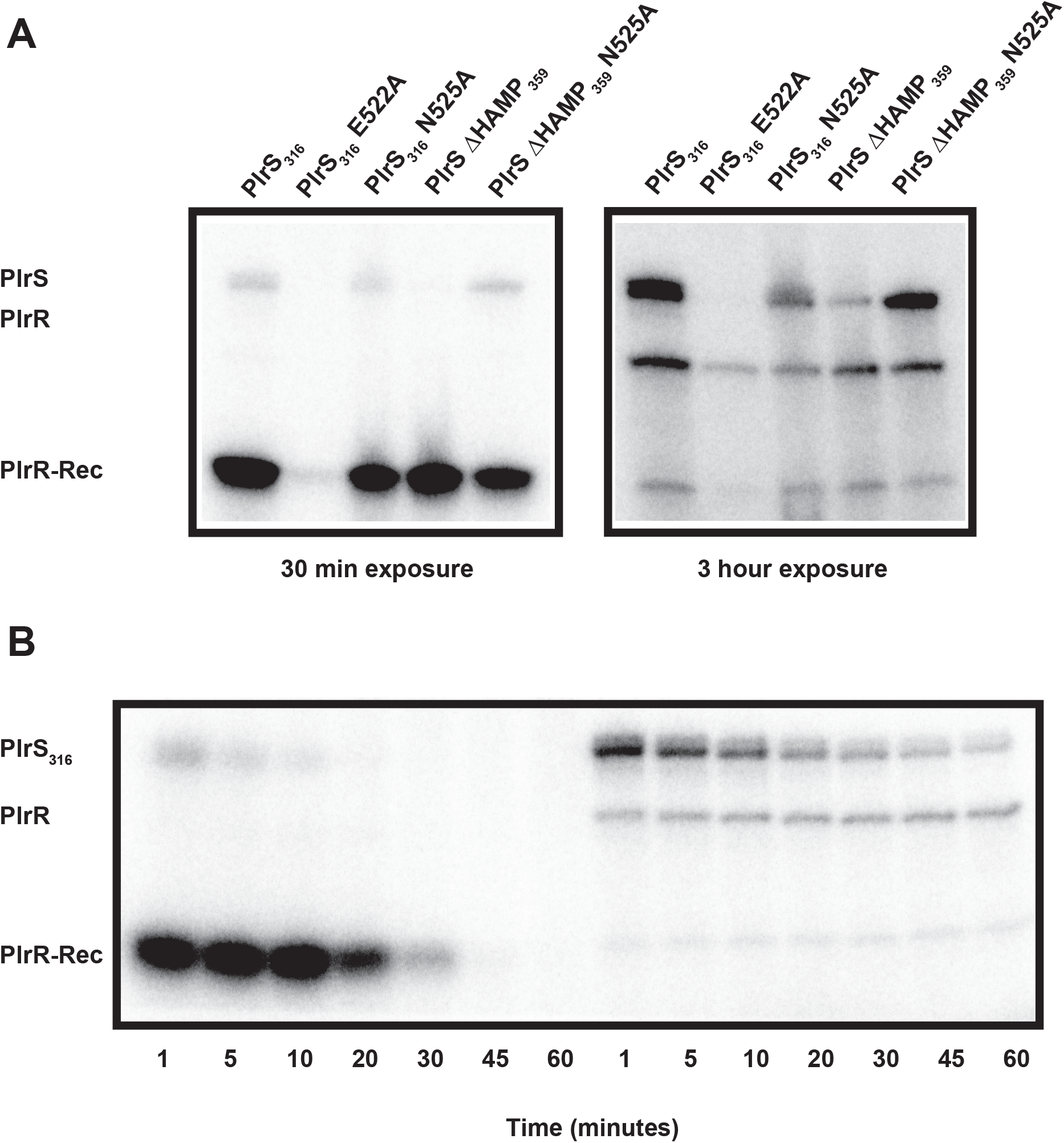
Phosphotransfer from PlrS to PlrR and PlrR-Rec. A. PlrS proteins were allowed to autophosphorylate with [γ-^32^P]ATP and then mixed with PlrR (right) or PlrR-Rec (left) at a 1:1 molar ratio to allow for phosphotransfer over 10 minutes at room temperature. B. PlrS_316_ autophosphorylated and then mixed 1:1 with PlrR (left) or PlrR-Rec (right) over 1 hour at room temperature. Experiments were performed in triplicate. Representative phosphorimages are shown.

### The phosphoryl group on PlrR is relatively stable

To assess the stability of phosphoryl groups on PlrR, we generated PlrR-P. As previously noted, PlrR-P could not be generated using small molecule phosphodonors, therefore requiring phosphorylation by a donor kinase (*i.e*., PlrS). However, various schemes to separate full-length PlrR from PlrS were unsuccessful. Therefore, to measure autodephosphorylation of full-length PlrR in the presence of PlrS we (i) blocked PlrS phosphatase activity with the N525A substitution (see later section) and (ii) selectively diminished phosphotransfer from PlrS to PlrR. The PlrS His phosphorylation site must be unprotonated to autophosphorylate with ATP and protonated to serve as a phosphodonor for PlrR (Kennedy *et al*., 2022; Mayover *et al*., 1999). Following incubation of PlrS ΔHAMP_359_ N525A with PlrR and [γ-^32^P]ATP, we raised the pH by 3.2 units, which reduced the rate of phosphotransfer from PlrS to PlrR by 10^3.2^ (1600 fold), allowing autodephosphorylation to occur without significant replenishment of PlrR-P (Figure 6A). In contrast to reactions involving His residues, autodephosphorylation kinetics of response regulator Asp residues are pH independent (Silversmith *et al*., 1997). k_dephos_ of PlrR was 0.21 ± 0.02 hr^−1^ (n = 4).

**Figure 6.**
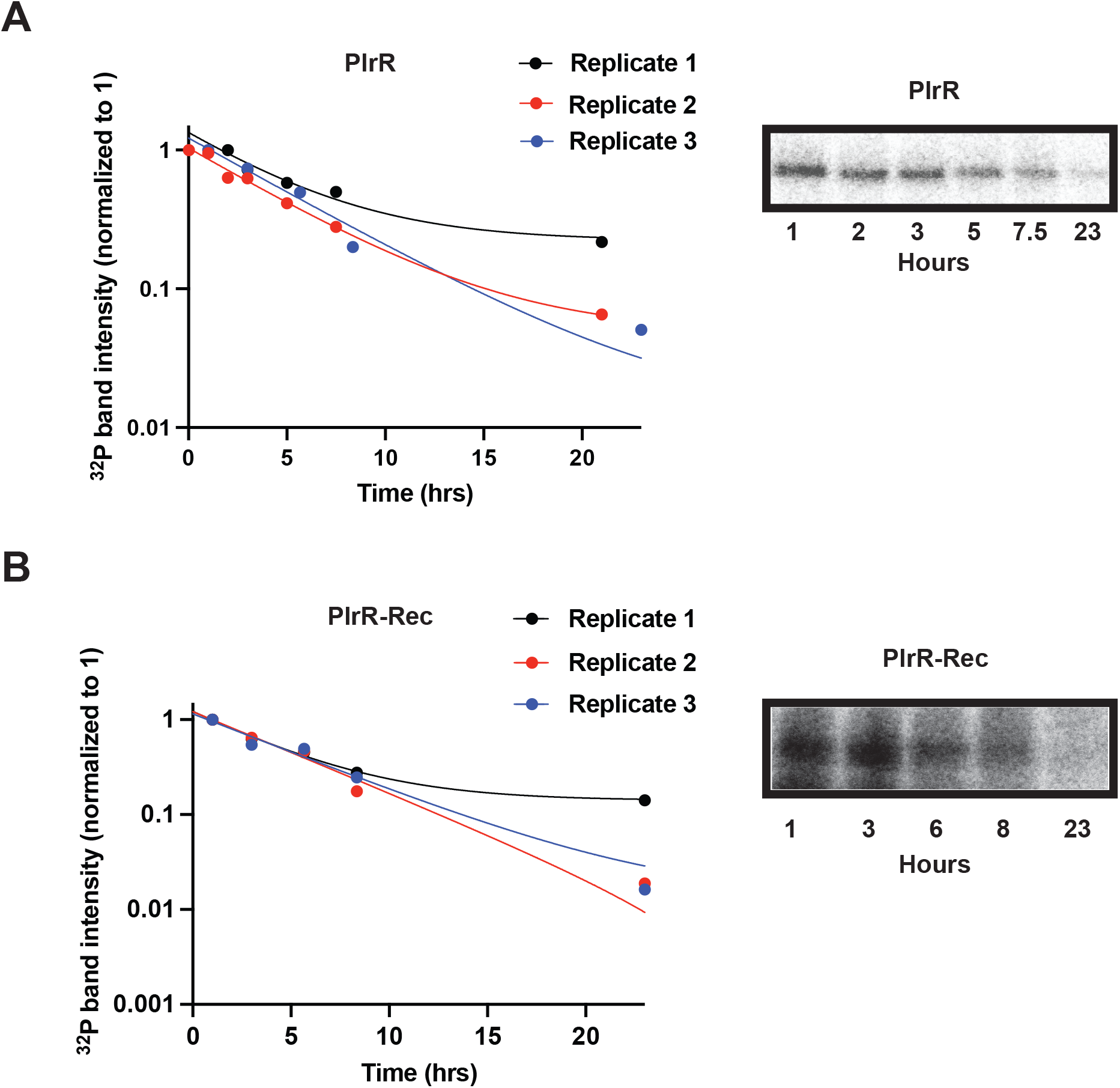
PlrR autodephosphorylation kinetics. Representative phosphorimages (right) and data plots (left) of (A) PlrR and (B) PlrR-Rec autodephosphorylation experiments. Autodephosphorylation was conducted at room temperature over 23 hours to measure rate constants. PlrS ΔHAMP359 N525A and [γ-^32^P]ATP were used to phosphorylate PlrR or PlrR-Rec for 2 hours. Then pH was raised 3.2 units to reduce phosphotransfer. Time points were taken after pH was raised. For all experiments, data were normalized to the maximal value and fit to an exponential decay equation with a plateau reflecting background noise.

To assess the role, if any, of the DNA binding domain in autodephosphorylation, we generated PlrR-Rec-P and applied the same method as with PlrR-P (Figure 6B). k_dephos_ of PlrR-Rec was 0.24 ± 0.07 hr ^−1^ (n = 4), indistinguishable from PlrR, suggesting that the DNA binding domain did not influence the rate of autodephosphorylation of PlrR. When either PlrR or PlrR-Rec was mixed with [γ-^32^P]ATP alone, we did not observe any label accumulation (data not shown) demonstrating that the presence of residual [γ-^32^P]ATP in the experiment did not interfere with our ability to exclusively observe autodephosphorylation.

The ~3 hour half-life of the phosphoryl group on PlrR and PlrR-Rec suggests that signal depletion requires phosphatase activity from an external source (*e.g*., PlrS) and that the functional biological output of the PlrSR TCS (likely transcriptional regulation) works on a longer timescale than something like chemotaxis (CheB and CheY autodephosphorylation half-lives are on the order of tens of seconds (Thomas *et al*., 2008)).

### PlrS exhibits phosphatase activity toward PlrR

Many sensor histidine kinases exhibit phosphatase activity toward their partner response regulators to respond rapidly to abrupt changes in environmental conditions and turn the system off as quickly as the system is turned on via kinase activity. Additionally, phosphatase activity is critical to erase spurious crosstalk from other sensor kinases or non-specific autophosphorylation of response regulators by small molecule phosphodonors, which would decouple the signal from the stimulus (Huynh & Stewart, 2011). To determine if PlrS can dephosphorylate PlrR-P, we incubated 1 μM PlrS with 10 μM PlrR-Rec in the presence of 100 μM [γ-^32^P]ATP. We focused our attention on PlrR-Rec because the experiments of Figure 5 suggested PlrS exhibited much greater reactivity toward PlrR-Rec than full-length PlrR. As noted previously, we could not generate the PlrR-Rec-P substrate for the phosphatase reaction with small molecule phosphodonors, and for technical reasons we could not remove residual ATP from the reaction with PlrS after phosphotransfer, so we could not measure phosphatase activity directly. We instead inferred phosphatase activity from the product balance of multiple simultaneous reactions. Incubation of PlrS, PlrR-Rec, and ATP results in (i) autophosphorylation of PlrS with ATP, (ii) phosphotransfer from PlrS-P to PlrR-Rec, (iii) autodephosphorylation of PlrR-Rec-P with water, and (iv) potential phosphatase activity of PlrS toward PlrR-Rec-P. PlrS_316_ N525A should be defective for phosphatase activity because the Ala at H+4 cannot coordinate the attacking water molecule (Huynh *et al*., 2010; Huynh & Stewart, 2011; Mideros-Mora *et al*., 2020; Willett & Kirby, 2012). Thus, phosphatase activity can be inferred by comparing the level of PlrR-Rec-P in the presence of PlrS proteins bearing Asn or Ala at H+4. Radiolabel accumulation on PlrR-Rec was much slower in the presence of PlrS_316_ than PlrS_316_ N525A (Figure 7A, left and middle). Furthermore, radiolabel was detectable, albeit barely, on PlrS_316_ N525A but not PlrS_316_. The differences in radiolabel accumulation between PlrR-Rec and the corresponding kinase were as expected if PlrS_316_ had phosphatase activity but PlrS_316_ N525A did not.

**Figure 7.**
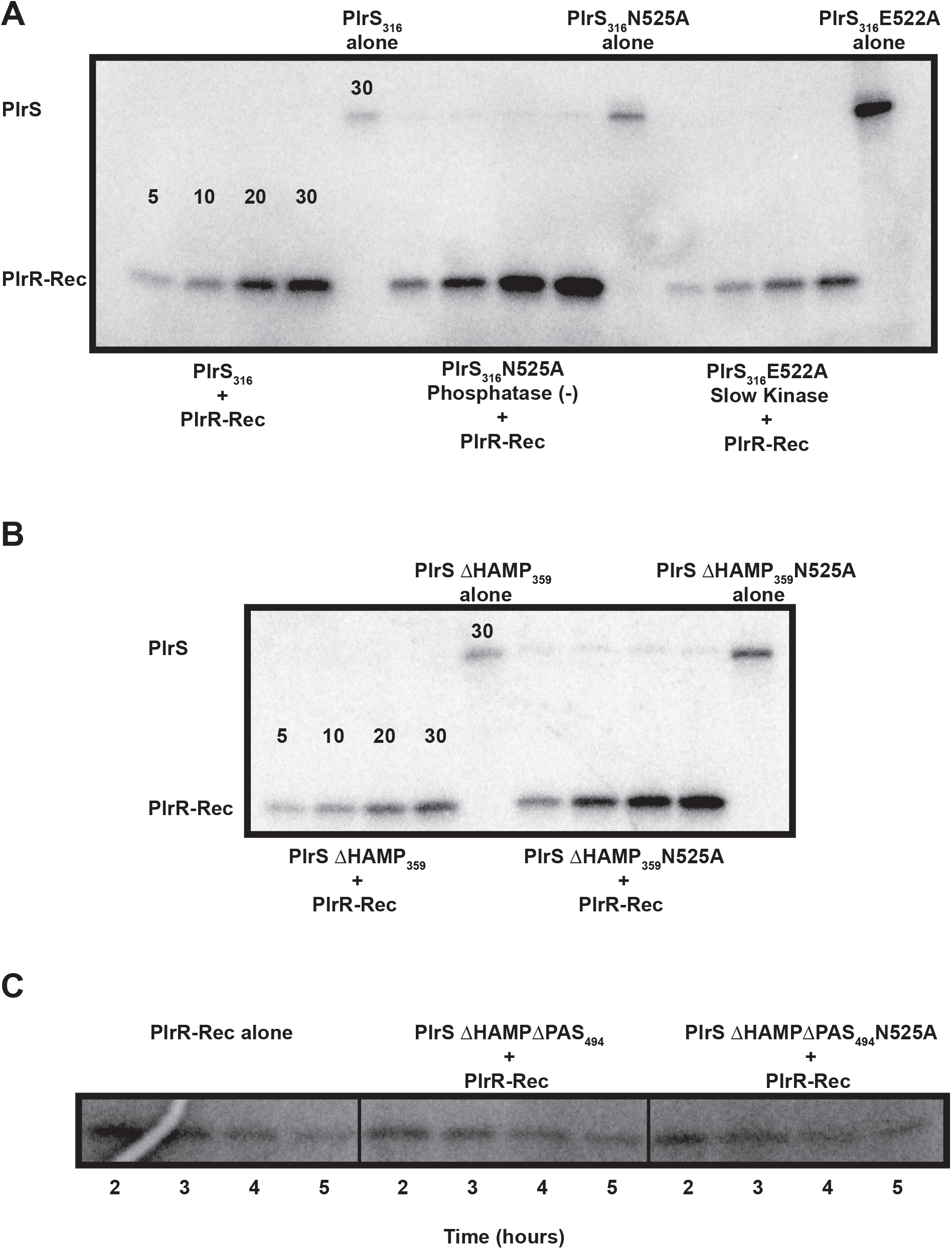
PlrS phosphatase activity toward PlrR-Rec. A. Representative phosphorimage of PlrR-Rec and [γ-^32^P]ATP mixed with PlrS_316_ (left), PlrS_316_ N525A (middle), or PlrS_316_ E522A (right) at a 10:1 ratio over the course of 30 minutes at room temperature. Lanes 5, 10 and 15 are PlrS_316_, PlrS_316_ N525A, PlrS_316_ E522A respectively at 5 μM mixed with [γ-^32^P]ATP for 30 minutes. B. Representative phosphorimage of PlrR-Rec and [γ-^32^P]ATP mixed with PlrS ΔHAMP_359_ (left) or PlrS ΔHAMP_359_ N525A (right) at a 10:1 ratio over the course of 30 minutes at room temperature. Lanes 5 and 10 are PlrS ΔHAMP_359_ (left) or PlrS ΔHAMP_359_ N525A (right) respectively at 5 μM mixed with [γ-^32^P]ATP for 30 minutes. C. PlrR-Rec-P alone (left), PlrR-Rec-P with PlrS ΔHAMPΔPAS_494_ (middle), and PlrR-Rec-P with PlrS ΔHAMPΔPAS_494_ N525A (right) at 10:1 ratios over 5 hours at room temperature.

PlrS_316_ E522A is defective for autophosphorylation (Figs. 2 & 3, Table 1), but contains the wild-type Asn525 residue, so should not be defective for phosphatase activity. Therefore, it is likely that the slower radiolabel accumulation on PlrR-Rec in the presence of PlrS_316_ E522A compared to PlrS_316_ (Figure 7A, left and right) is due to slower autophosphorylation of PlrS_316_ E522A rather than differences in phosphotransfer or phosphatase activity. PlrS_316_ H521Q did not exhibit detectable phosphatase activity toward PlrR-Rec (data not shown), despite containing Asn525. Due to the inability of PlrS_316_ H521Q to autophosphorylate, phosphatase activity was assessed by a different method than that shown in Figure 7, as described in Experimental Procedures. Although the conserved His residue of HisKA sensor kinases contribute to the phosphatase reaction (Mideros-Mora *et al*., 2020), there are many cases demonstrating that the His is not essential for phosphatase activity (Huynh & Stewart, 2011; Willett & Kirby, 2012), suggesting that Gln521 interfered.

### The PAS domain is critical for PlrS phosphatase activity toward PlrR

To assess whether the HAMP domain contributes to PlrS phosphatase activity, we compared the radiolabel accumulation of PlrR-Rec when mixed with PlrS ΔHAMP_359_ vs PlrS ΔHAMP_359_ N525A. The pattern of radiolabel accumulation on PlrR-Rec with PlrS ΔHAMP_359_ vs PlrS ΔHAMP_359_ N525A mimicked that of PlrS_316_ vs PlrS_316_ N525A, suggesting that the HAMP domain is not critical to phosphatase activity (Figure 7B). However, diminished accumulation of PlrR-Rec-P could also be due to reduced autophosphorylation of PlrS ΔHAMP_359_ compared to PlrS ΔHAMP_359_ N525A (Figure 3A).

Because autophosphorylation of PlrS ΔHAMPΔPAS_494_ and PlrS ΔHAMPΔPAS_494_ N525A and autodephosphorylation of PlrR-Rec occurred on similar time scales, we could assess phosphatase activity directly by adding either PlrS ΔHAMPΔPAS_494_ or PlrS ΔHAMPΔPAS_494_ N525A to radiolabeled PlrR-Rec. Neither PlrS ΔHAMPΔPAS_494_ nor PlrS ΔHAMPΔPAS_494_ N525A increased dephosphorylation of PlrR-Rec (Figure 7C). We also assayed potential phosphatase activity of the two longer PlrS ΔHAMPΔPAS constructs (PlrS ΔHAMPΔPAS_477_, PlrS ΔHAMPΔPAS_484_) but again failed to detect any (Figure S2). Collectively, these data suggest the PAS domain is critical for PlrS phosphatase activity.

### *PlrS phosphatase activity is required for* B. bronchiseptica *persistence in the murine lower respiratory tract*

Strains predicted to lack PlrS kinase activity are unable to persist in the LRT (Bone *et al*., 2017; Kaut *et al*., 2011). To determine if PlrS phosphatase activity is also required for bacterial survival in the LRT, we constructed a strain that, based on our *in vitro* phosphorylation assays, should produce a PlrS protein that retains kinase activity, but lacks phosphatase activity. *plrR* appears to be essential, and plasmid insertions near the 3’ end of *plrS* are not tolerated (Kaut *et al*., 2011), suggesting the presence of an essential promoter for *plrR* within *plrS*. Therefore, to construct the desired stain, we first placed *plrR*, driven by the constitutively active S12 promoter, at the att*Tn7* site on the chromosome. We then deleted *plrR* from its native site to create a strain called Δ*plrR att*Tn7::*plrR*. We then constructed one derivative of this strain with an in-frame deletion removing the codons for PlrS amino acids 5 through 198 (Figure 1) (*plrS*Δ_5-198_ Δ*plrR att*Tn7::*plrR*) and another encoding PlrS with the N525A substitution (*plrS N525A* Δ*plrR att*Tn7::*plrR*). All strains grew at similar rates *in vitro* (data not shown).

We inoculated BALB/c mice intranasally with 7.5 x10^4^ colony forming units (CFUs) of the four strains and quantified CFUs recovered from the nasal cavity and lungs at three hours, two days, and seven days post-inoculation (Figure 8). Similar numbers of CFUs of the wild-type and Δ*plrR att*Tn7::*plrR* strains were recovered from nasal cavities and lungs at every time point, indicating that constitutive expression of *plrR in trans* was sufficient for, and not detrimental to, bacterial survival during infection. Consistent with our previous data using a *plrS*Δ_5-198_ strain (Bone *et al*., 2017; Kaut *et al*., 2011), the *plrS*Δ_5-198_ Δ*plrR att*Tn7::*plrR* strain persisted in the nasal cavity similarly to the wild-type strain, but was cleared, or nearly cleared, from the lungs by day two post-inoculation. The *plrS N525A* Δ*plrR att*Tn7::*plrR* strain was similarly able to persist in the nasal cavity, but was severely defective for persistence in the lungs compared to the wild-type and Δ*plrR att*Tn7::*plrR* strains, although it was recovered at approximately two to three logs higher at two and seven days post-inoculation than the *plrS*Δ_5-198_ Δ*plrR att*Tn7::*plrR* strain. Together, these data indicate that both kinase and phosphatase activity of PlrS are required for persistence of *B. bronchiseptica* in the LRT.

**Figure 8.**
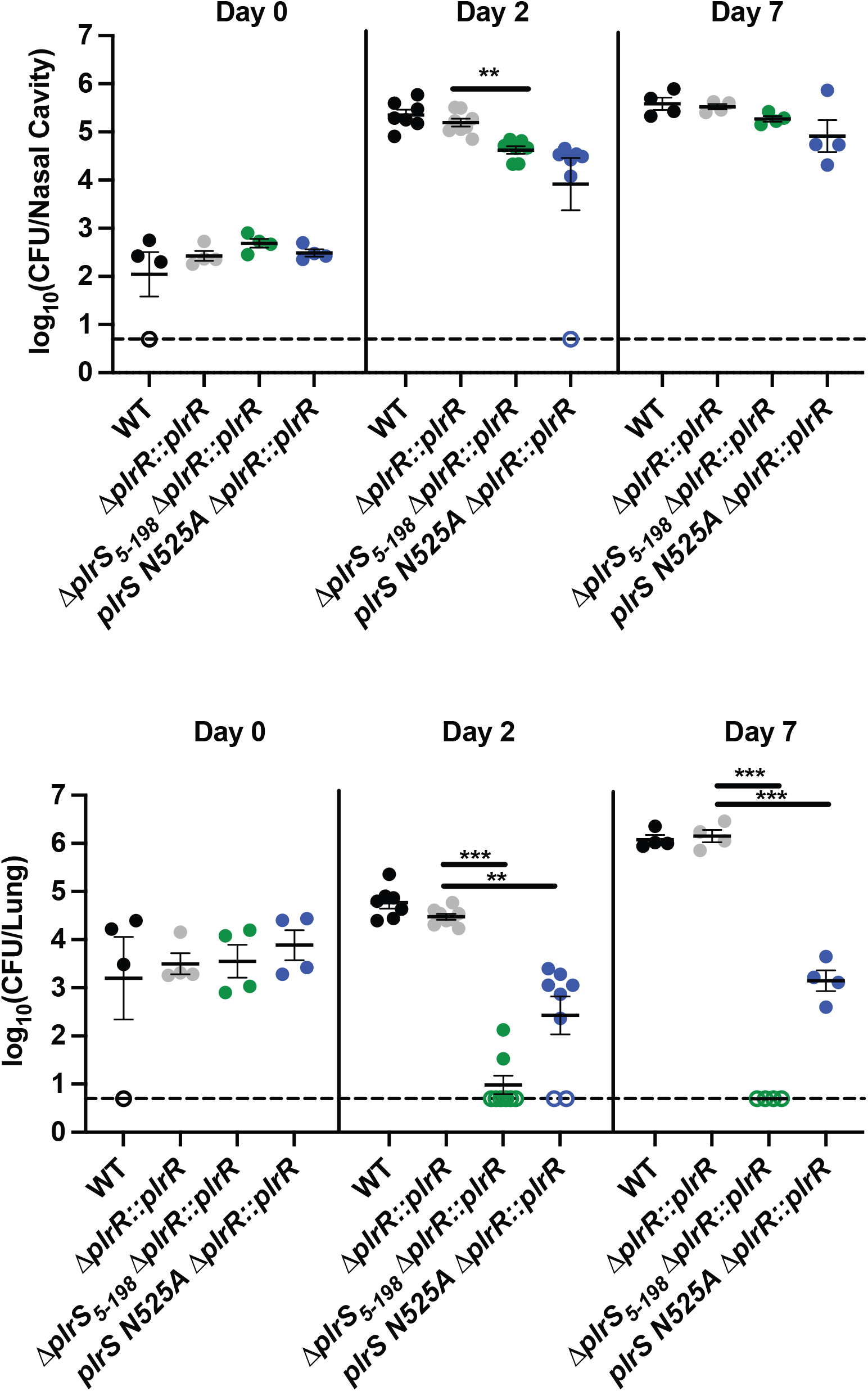
PlrS phosphatase activity is required for persistence *of B. bronchiseptica* in the LRT of mice. CFU recovered from the nasal cavities and lungs of BALB/c mice infected intranasally with wild-type and mutant *B. bronchiseptica* strains are shown. Each strain with the designation Δ*plrR::plrR* has been abbreviated to represent Δ*plrR att*Tn7::*plrR*. Each spot represents CFU recovered from a single animal. Dashed lines indicate the lower limit of detection, and points outlined with no fill color represent samples from which no CFU were recovered. Horizontal bars represent the mean and whiskers show the standard error of the mean. Statistical significance was determined by unpaired, two-tailed T-tests with *, **, *** representing p<0.01, p<0.001, and p<0.0001, respectively.

## 6 DISCUSSION

PlrS and PlrR are critical to the survival of *B. bronchiseptica* in the LRT (Bone *et al*., 2017; Kaut *et al*., 2011). Our study used phosphorylation chemistry to establish that PlrS and PlrR behave as a functional two-component system. We showed that PlrS can exhibit both kinase and phosphatase activity towards PlrR, and we identified amino acids that are critical for each respective activity. PlrR is an unusual response regulator in which the DNA binding domain inhibits phosphotransfer and phosphatase activity by PlrS and facilitates tetramer formation. We constructed a *B. bronchiseptica* mutant that should, based on our biochemical data, retain PlrS kinase activity but lack phosphatase activity. The PlrS phosphatase-defective *B. bronchiseptica* mutant was severely defective for persistence in the LRT of mice, demonstrating the importance of PlrS phosphatase activity *in vivo*. Altogether, the data presented in this study support a phosphorylation dependent virulence model wherein PlrS responds to stimuli present in the host by adjusting its kinase and phosphatase activities to control PlrR-P levels to regulate gene expression patterns that promote bacterial survival in the LRT.

### The biochemistry of PlrS

PlrS and PlrR share amino acid sequence similarity with NtrY/NtrX-like histidine sensor kinases and response regulators (Kaut *et al*., 2011). NtrY proteins are characterized by multiple transmembrane segments followed by cytoplasmic HAMP, PAS, and HisKA domains, as is the case for PlrS. The NtrYX system was first characterized as important for nitrogen metabolism in *Azorhizobium caulinodans* but has been identified and biochemically explored in other pathogenic alphaproteobacteria such as *Brucella abortis* and *Caulobacter crescentus* (Carrica Mdel *et al*., 2012; Pawlowski *et al*., 1991; Stein *et al*., 2021). To the best of our knowledge, NtrY-like proteins in betaproteobacteria have not previously been shown to function as canonical sensor histidine kinases.

The PlrS H521Q protein did not exhibit autophosphorylation, consistent with the expectation that His521 is the only site of phosphorylation in PlrS. Furthermore, PlrS H521Q did not exhibit phosphatase activity toward PlrR, consistent with the expectation that His521 contributes to PlrS phosphatase activity (Mideros-Mora *et al*., 2020). We also established that two amino acids adjacent to His521, Glu522, and Asn525, behaved similarly in phosphorylation chemistry relative to the analogous positions in other sensor kinases (Atkinson & Ninfa, 1993; Dutta *et al*., 2000; Hsing *et al*., 1998; Pazy *et al*., 2010; Silversmith, 2010; Willett & Kirby, 2012). Glu522 was essential to maintain a high rate of autophosphorylation whereas Asn525 was essential for facilitating phosphatase activity even in the context of His521. Although NtrY phosphatase activity has been identified in alphaproteobacteria (Stein *et al*., 2021), we demonstrated the importance of PlrS phosphatase activity for virulence of *B. bronchiseptica*, a betaproteobacterium.

Although neither kinase nor phosphatase activities were dictated by the HAMP domain, PlrS proteins lacking the PAS domain were deficient in both. However, the negative data does not suggest how the PAS domain might exert its influence on the biochemical activity of PlrS and leaves us to speculate. A *Streptococcus pneumoniae* truncated WalK sensor kinase lacking the PAS domain was deficient for phosphatase activity, but still maintained autophosphorylation (Gutu *et al*., 2010), contrasting with *B. subtilis* KinA, which relies upon the PAS domain for kinase activity (Lee *et al*., 2008). Although the stimuli detected by PlrS have not been identified, the importance to persistence in the lung suggests that stimuli might have to do with O2, CO2, or redox levels, all of which would be consistent with sensing by a PAS domain as evidenced by other NtrYX systems (Atack *et al*., 2013; Carrica Mdel *et al*., 2012). We expect that the PAS domain stabilizes both kinase and phosphatase conformations depending on ligand binding. Stimuli detection by the periplasmic PDC domain might also affect the balance of kinase and phosphatase activities via conformational changes transmitted through the PAS domain.

### PlrR is an unusual response regulator

Some NtrX homologs in betaproteobacteria (including PlrR) lack the central AAA+ ATPase domain found in NtrX proteins of alphaproteobacteria (Atack *et al*., 2013; Bonato *et al*., 2016). We observed additional differences from previously characterized NtrX proteins. *B. abortis* NtrX autophosphorylates with small molecule phosphodonors (Fernandez *et al*., 2015), whereas PlrR did not. This may represent a biochemical functional difference between NtrX proteins of alpha- and betaproteobacteria. As expected for the NtrC family (Gao *et al*., 2019), *B. abortis* NtrX exhibits a monomer-dimer equilibrium and forms dimers using the α4α5 interface (Fernandez *et al*., 2015), whereas PlrR formed a dimer-tetramer equilibrium. This may represent a structural/functional difference between NtrX proteins of the alpha- and betaproteobacteria.

More commonly, response regulators exist as a population balance between monomers and dimers, depending on the active/inactive conformational state. Phosphorylation stabilizes active conformations, which results in dimerization (Gao *et al*., 2019). The dimeric state is often associated with being phosphorylated and facilitates output function. Many response regulators contain DNA binding output domains that bind to dimeric DNA sequences when dimerized and activate transcription. It is often presumed that an unphosphorylated RR is functionally inactive while a phosphorylated RR is functionally active because phosphorylation and dimerization tend to follow suit.

Although we cannot yet determine if either dimeric or tetrameric PlrR act as transcriptional activators or repressors or even bind DNA at all, the strong homology between PlrR and the transcriptional response regulator NtrX leads us to predict PlrR will bind DNA and regulate gene expression. A tetramer oligomeric state is unusual for a response regulator but not unprecedented. In *Shewanella oneidensis*, the response regulator HnoC exists as a tetramer when unphosphorylated (Plate & Marletta, 2013). Upon phosphorylation, HnoC dissociates into a dimer and in turn has a lower affinity for its promoters, leading to derepression. In *Pseudomonas aeruginosa*, the diguanylate cyclase response regulator WspR requires tetramerization to be active (De *et al*., 2009) and the phosphodiesterase response regulator RocR forms an asymmetrical tetramer where the AB side restricts ligand access to the active site of the CD side (Chen *et al*., 2012). We observed that the DNA binding domain of PlrR inhibited both phosphotransfer and phosphatase activity of PlrS. It is possible that the PlrR tetramer structure, like RocR, prevents PlrS from efficient interaction. In contrast to PlrR, the response regulator RR_1586 in *Clostridiodes difficile* tetramerizes upon phosphorylation (Hebdon *et al*., 2018). However, RR_1586 phosphorylation (and therefore tetramerization) decreases the affinity for target DNA similarly to HnoC. In the context of many examples with many unique mechanisms, it is difficult to predict whether PlrR is active as a tetramer, dimer, or both. We expect that PlrR exhibits differential regulation as a tetramer and a dimer by acting as repressor at some sites and/or activator at others. Establishing the exact activity of each PlrR oligomeric state will require additional investigation.

### *The balance between PlrS kinase and phosphatase activities* in vivo

The observations that Δ*plrS* and *plrS H521Q* strains were cleared quickly from the LRT led to the conclusion that PlrS kinase activity is important for persistence (Bone *et al*., 2017; Kaut *et al*., 2011). We showed in this study that PlrS phosphatase activity is also important for persistence in the lung. It is not known if PlrSR regulates gene expression when PlrR is phosphorylated or unphosphorylated. Based upon the dual importance of both PlrS kinase and phosphatase activity, we propose that PlrR and PlrR-P may have different regulatory activities that require a fine balance between kinase and phosphatase activity from PlrS. Because our new data show that a PlrS H521Q protein is deficient for both kinase and phosphatase activity, our previous conclusion that PlrS kinase activity is required *in vivo* based solely on this protein is not justified. However, the different and more severe phenotype of *plrS H521Q*, in rats, (Kaut et al., 2011) compared to *plrS N525A*, in mice, strongly suggests that PlrS kinase activity is required *in vivo* (unless there is a subtle difference between rodent models).

If only phosphorylated PlrR was essential, then we would not have expected such a severe defect of the *plrS N525A* mutant within the host. There are other examples to support the notion that the timing and concentration of both the phosphorylated and unphosphorylated forms of a response regulator are critical for survival. *S. pneumoniae* is attenuated for virulence in a pneumonia model when the WalK sensor histidine kinase is defective for phosphatase activity (Gutu *et al*., 2010). In group A *Streptococcus*, the phosphatase activity of the CovS sensor histidine kinase is critical for establishing skin infections and surviving neutrophil attack. Without the ability to be dephosphorylated, the CovR response regulator constitutively represses critical genes for survival (Horstmann et al., 2018). Similarly, *P. aeruginosa* relies upon KinB phosphatase activity towards its partner response regulator AlgB, which when phosphorylated, represses critical genes for virulence (Chand *et al*., 2012).

Our data does not elucidate the exact balance between kinase and phosphatase activities of PlrS. A histidine kinase may have multiple conformations that allow for kinase activity and multiple conformations that allow for phosphatase activity, but each activity is mutually exclusive and is determined by ligand binding/dissociation-induced conformational changes (Buschiazzo & Trajtenberg, 2019; Jacob-Dubuisson *et al*., 2018). It is possible that PlrS has coevolved with PlrR to have very stable kinase and phosphatase conformations to account for the structural obstruction from the PlrR DNA binding domain.

### The search for a PlrS stimulus and PlrR regulon

While our data contributes to understanding how PlrS and PlrR function within *Bordetella*, there are still gaping holes. Three key questions remain: (i) what stimuli regulate PlrS activity? (ii) what genes are regulated by PlrR? and (iii) how are the BvgAS (previously thought of as the master virulence regulator) and PlrSR TCSs functionally connected? The results reported here do not directly address these key questions, but provide some insights on mechanisms (*e.g*., PlrR forms tetramers) and may supply characterized PlrS mutant proteins (H521Q and N525A) that could help unravel the answers. Identifying the oligomeric states in PlrR will be fruitful for DNA binding experiments to identify target loci. Identifying the importance of the PAS domain will spur targeted mutagenesis and a smaller pool of stimuli to test for PlrS. Eventual understanding of the PlrSR TCS could help identify new vaccine and anti-virulence targets for harmful *Bordetella* species.

## 7 EXPERIMENTAL PROCEDURES

### Media conditions

All *E. coli* strains used in this study were grown in Lysogeny Broth (LB) (10 g tryptone, 5 g yeast extract, 10 g NaCl per liter) with the antibiotic kanamycin (Km; 30 or 50 μg ml^−1^) as noted.

All *B. bronchiseptica* strains were grown on Bordet-Gengou (BG) agar supplemented with 6% defibrinated sheep blood (Hemostat, catalog no. DSB1) or in Stainer-Scholte (SS) broth supplemented with SS supplement at 37° C (Stainer & Scholte, 1970). As needed, media were supplemented with streptomycin (Sm; 20 μg ml^−1^), gentamicin (Gm; 30 μg ml^−1^), kanamycin (Km; 125 μg ml^−1^), or diaminopimelic acid (DAP; 300 μg ml^−1^).

### Strain construction

Descriptions of each strain are given in Table S1. Relevant plasmids are listed in Table S2 with corresponding oligonucleotides listed in Table S3.

The pET28a (+) plasmids used for production of His_6_ fusion proteins were created for this study. Plasmids pET28a (+) PlrS_316_, pET28a (+) PlrS ΔHAMP_359_, pET28a (+) PlrS ΔHAMPΔPAS_494_, pET28a (+) PlrS ΔHAMPΔPAS 477, pET28a (+) PlrS ΔHAMPΔPAS_484_, pET28a (+) PlrR, and pET28a (+) PlrR-Rec listed in Table S1 were synthesized by Genewiz using codons optimized for protein production in *E. coli*. To generate point mutations in each of the parent plasmids, we used Platinum Superfi Polymerase in a site directed mutagenesis PCR reaction using the primers in Table S3. Mutants were confirmed by sequencing. We used the NEB5*α* high efficiency strain for plasmid propagation and the Invitrogen BL21 Star cells for protein production and purification. Kanamycin (30 μg ml^−1^) was used for propagation of protein producing strains.

We used the DH5α *E. coli* strain (supplemented with Km 50 μg ml^−1^) for plasmid construction and propagation of plasmids needed to construct *B. bronchiseptica plrSR* mutant strains. Any mutations made within *B. bronchiseptica* strains were confirmed by PCR and/or sequencing. The Δ*plrR att*Tn7::*plrR* containing a constitutively expressed *plrR* at an exogenous location and the deletion of *plrR* at the native locus was generated by first introducing *plrR* at the att*Tn7* the pUCS12-*plrR* and pTNS_3_ plasmids, then deleting the native *plrR* using pMAB09 by allelic exchange as previously described (Inatsuka *et al*., 2010). The Δ*plrR att*Tn7::*plrR* strain was then used to generate further strains using allelic exchange: Δ*plrS_5-198_*Δ*plrR att*Tn7::*plrR* using pMD11 and *plrS N525A* Δ*plrR att*Tn7::*plrR* using *plrS N525A*::pEG7

### Protein expression and purification

Each BL21 strain was inoculated in 5 mL of LB Km to grow at 37 °C overnight. Overnight cultures were inoculated into 1 L of LB Km to grow at 37 °C until the culture reached the appropriate OD_600_. For proteins below 30 kDa, the cultures were grown to between 0.6-0.8 OD_600_ and induced with 1 mM of IPTG for 20 hours at room temperature. For proteins above 30 kDa, the cultures were grown to between 0.06-0.1 OD_600_ and induced with 250 μM of IPTG for 20 hours at room temperature. Cultures were collected by centrifugation at 4200 × *g* for 30 mins at room temperature. Supernatants were discarded and cell pellets were resuspended in 25 ml lysis buffer (50 mM Hepes pH 7.0, 50 mM KCl, 150 mM NaCl, 10 mM MgCl_2_, 10% (v/v) glycerol) and frozen. Frozen pellets were thawed and lysed using an Avestin Emulsiflex C3 homogenizer. The lysate was separated from cell debris by centrifugation at 145,000 × *g* for 45 mins at 4 °C. His-tagged proteins were then captured using Ni-NTA resin. Nonspecific protein was removed using sequential washes (20 mL of 40 mM and 20 mL of 80 mM imidazole) and his-tagged protein was eluted using 150 mM and 300 mM imidazole. Proteins were then dialyzed to remove imidazole before experiments.

### Radioisotope use and detection

For every experiment using [γ-^32^P]ATP, we used a 1:20 mix of 10 μCi (6,000 Ci/mmol; PerkinElmer) and unlabeled ATP (100 μM unless otherwise noted). All radioactive protein samples were separated by size on a 4-to-20% Mini-Protean TGX precast SDS-PAGE gel (Bio-Rad). The gel was then dried and imaged using a Typhoon phosphorimager (Molecular Dynamics GE Healthcare Life Sciences). All biochemical experiments were conducted in at least triplicate unless otherwise noted.

### Autophosphorylation of PlrS constructs

Protein concentration was estimated using the A_280_ and extinction coefficients [PlrS_316_ (23950 M^−1^ cm^−1^), PlrS ΔHAMP_359_ (23950 M^−1^ cm^−1^), PlrS ΔHAMPΔPAS_494_ (15470 M^−1^ cm^−1^), PlrS ΔHAMPΔPAS_484_ (16960 M^−1^ cm^−1^), PlrS ΔHAMPΔPAS_477_ (16960 M^−1^ cm^−1^), PlrR (16960 M^−1^ cm^−1^), PlrR-Rec (12490 M^−1^ cm^−1^) calculated by ExPASy ProtParam tool (Gasteiger, 2005)]. Autophosphorylation reactions were initiated by the addition of 100 μM [γ-^32^P]ATP to protein (either 1 or 10 μM) in reaction buffer (50 mM HEPES pH 7.0, 50 mM KCl, 10 mM MgCl_2_). At specific timepoints, sample was removed, and the reaction stopped by the addition of Laemmli sample buffer (0.25 M Tris-HCl pH 6.8, 10% [w/v] SDS, 50% [v/v] glycerol, 0.005% [w/v] bromophenol blue).

For the single time point autophosphorylation reactions in Figure 2, each PlrS construct was allowed to incubate with [γ-^32^P]ATP for 15 minutes. Rate constants for autophosphorylation were determined over the course of an hour on ice (0 °C). We used ImageJ software to measure the pixel density of each time point and normalized to the highest pixel density band for each experiment. GraphPad Prism software was used to fit the data to an exponential decay model to determine the rate constant at a single concentration of ATP.

### Fluorescence intensity measurements

PlrR encodes a tryptophan residue located two residues C-terminal to the predicted Asp52 phosphorylation site. In an attempt to measure autophosphorylation and/or autodephosphorylation rate constants using Trp fluorescence intensity, a PerkinElmer LS-50B fluorimeter was used to monitor fluorescence intensity changes. PerkinElmer FL WINLAB V.1.1 software was used to operate the instrument. All experiments were performed at 25 °C with constant stirring using an excitation wavelength of 295 nm and an emission wavelength of 346 nm. Autophosphorylation measurements were attempted using multiple small-molecule phosphodonors: phosphoramidate, monophosphoimidazole, or acetyl phosphate. A mixture of 10 mM MgCl_2_, 100 mM HEPES (pH 7.0), and 100 mM of each small-molecule phosphodonor were titrated separately using a Hamilton syringe into a reaction mixture of 5 mM PlrR, 10 mM MgCl_2_, 100 mM HEPES (pH 7.0), and 100 mM KCl in a quartz cuvette. No visible changes in fluorescence intensity could be observed that were not a direct result of dilution about each volume addition.

### Mass photometry

Concentration of protein for mass photometry is arbitrary. PlrR or PlrR D52A was diluted in PBS until the concentration was such that individual protein molecule impact events could be visualized on the Refeyn One Mass Photometer. Once the sample was appropriately diluted, impact and detachment events were recorded for 1 minute. The counted impact and detachments within 1 minute were considered a biological replicate. A BeF_3_^−^ solution (final 2 mM BeSO_4_ and 20 mM NaF) was added directly to protein sample and allowed to equilibrate before recording impacts and detachment events.

### Autodephosphorylation of PlrR and PlrR-Rec

All reactions were performed at room temperature in reaction buffer (50 mM HEPES pH 7.0, 50 mM KCl, 10 mM MgCl_2_). Phosphorylation of PlrR and PlrR-Rec was achieved by incubating 15 μM of either PlrR or PlrR-Rec with 1 μM PlrS ΔHAMP_359_ N525A in the presence of 15 μM [γ-^32^P]ATP mix. The reaction was allowed to proceed for 2 hours at room temperature. In order prevent continued phosphorylation of the response regulator to isolate only the dephosphorylation reaction a previously published pH jump method was used (Kennedy *et al*., 2022; Mayover *et al*., 1999). After initial phosphorylation of PlrR or PlrR-Rec, each reaction was combined with an equal volume of 500 mM sodium bicarbonate pH 10.5 for a final 200 mM sodium bicarbonate at a pH of 10.2. Following addition of the sodium bicarbonate samples were taken at various time points and quenched with stop buffer. Band intensities on each individual phosphorimage were quantified using ImageJ. GraphPad Prism software was used to fit the data to an exponential decay function with a plateau.

### Phosphotransfer experiments

For single time point phosphotransfer experiments (Figure 5A), 10 μM of PlrS constructs were mixed with 100 μM [γ-^32^P]ATP and allowed to autophosphorylate for approximately 18 minutes at room temperature. Then either PlrR or PlrR-Rec protein were added to a final concentration of 10 μM and phosphotransfer was terminated after 10 minutes by the addition of Laemelli sample buffer.

For phosphotransfer timecourse experiments (Figure 5B), 10 μM PlrS_316_ was mixed with were mixed with 100 μM [γ-^32^P]ATP and allowed to autophosphorylate for 15 minutes. Then either PlrR or PlrR-Rec were added to a final concentration of 10 μM, sample was removed, and reaction halted by addition of Laemelli sample buffer at specific time points over an hour at room temperature.

### Phosphatase experiments

1 μM PlrS construct was mixed with 10 μM PlrR-Rec and 100 μM [γ-^32^P]ATP. Sample was removed and reaction halted by addition of Laemelli sample buffer at specific time points over 30 minutes at room temperature. For each PlrS construct, we conducted an autophosphorylation reaction at 5 μM protein and 100 μM ATP for 30 minutes to confirm kinase function.

For PlrS ΔHAMPΔPAS_494_, PlrS ΔHAMPΔPAS_484_, and PlrS ΔHAMPΔPAS_477_ phosphatase experiments, 10 μM PlrR-Rec-P was generated using 3 μM PlrS_316_ N525A and 100 μM [γ-^32^P]ATP for 15 minutes. PlrR-Rec-P was then separated from PlrS_316_ N525A by passage through an Amicon 0.5 mL 50 kDa molecular weight cut off filter for 8 minutes at 16,000 x g. The flowthrough was then mixed with 1 μM ΔHAMPΔPAS_494_, PlrS ΔHAMPΔPAS_484_, or PlrS ΔHAMPΔPAS_477_ respectively. Sample was removed and reaction halted by addition of Laemelli sample buffer at specific time points over 30 minutes or 5 hours at room temperature.

### Phosphatase experiments for H521Q

10 μM PlrR-Rec-P was generated using 1 μM PlrS_316_ and 100 μM [γ-^32^P]ATP for 5 minutes. PlrR-Rec-P was then separated from PlrS_316_ by passage through an Amicon 0.5 mL 50 kDa molecular weight cut off filter for 8 minutes at 16,000 x g. The flowthrough was then mixed with 1 μM of PlrS H521Q or buffer alone. Sample was removed and reaction halted by addition of Laemelli sample buffer at specific time points over 2 hours at room temperature.

### Kinetic modeling of PlrS to PlrR phosphotransfer system

Data for phosphotransfer kinetics was simulated using the software KinTek Explorer (version 10.2.0, KinTek Corp.). Model inputs are listed below.

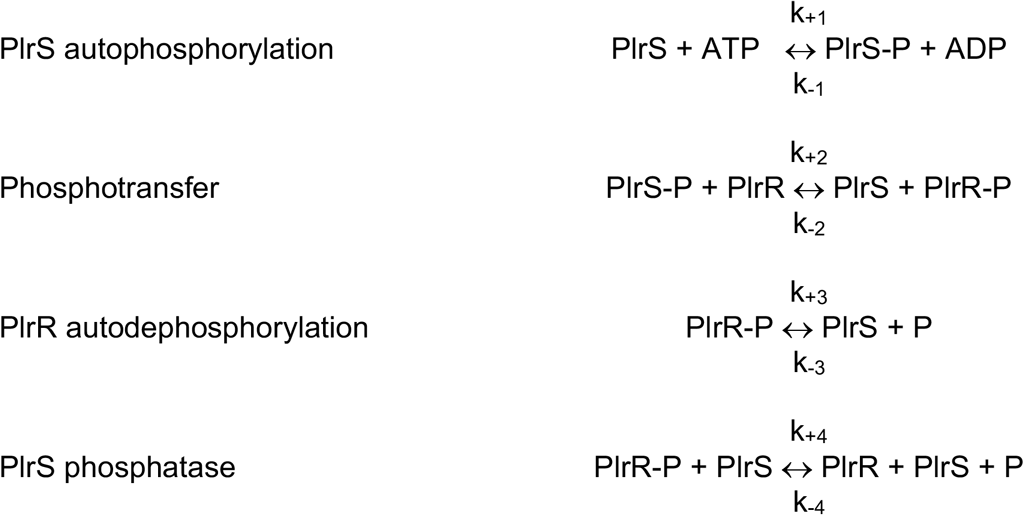

Bands from phosphotransfer time course experiments as in Figure 5B were quantified for both PlrS-P and PlrR-P or PlrR-Rec-P using ImageJ. These data were then set as observable outputs for PlrS-P and PlrR-P or PlrR-Rec-P. For the purposes of our analysis, rate constants for all reverse reactions (k_-1_, k_-2_, k_-3_, k_-4_) were fixed at zero. The initial autophosphorylation rate constant for PlrS was constrained to the experimentally determined value of 0.62 ± 0.4 min^−1^ (Table1). All other variables were allowed to be determined by the modeling software for simulations of experiments using PlrR-Rec. Variable k_+3_ obtained from the simulation for PlrR-Rec was then applied to data obtained with PlrR, *i.e*., PlrS autophosphorylation (k_+1_) and PlrR autodephosphorylation (k_+3_) were constrained.

### B. bronchiseptica *growth curve*

Cultures of *B. bronchiseptica* were grown for 18 hours at 37 °C in SS media supplemented with SS supplement and Sm. Each culture was normalized to 0.1 OD_600_/mL and grown on an orbital shaker at 37 °C. We measured the OD_600_ over 48 hours to identify the time of lag, exponential, and stationary phases. The doubling time was calculated during exponential phase growth between 0 and 8 hours using the equation t_D_= ln 2/k, where k = ln (OD_8hr_/OD_0hr_)/(8 hr – 0 hr) and t_D_ is doubling time.

### Bacterial colonization of the mouse respiratory tract

Six-week-old female BALB/c mice from Charles River Laboratories (catalog no. BALB/cAnNCrl) were inoculated intranasally with 7.5 × 10^4^ CFU *B. bronchiseptica* in 50 μL of Dulbecco PBS (2.7 mM KCl, 1.5 mM KH_2_PO_4_, 138 mM NaCl, 8 mM Na_2_HPO_4-_(H_2_O)_7_). Samples were collected at three hours, two days, and seven days post infection based on previous experiments using Δ*plrS_5-198_* (Bone *et al*., 2017). At each indicated time point, right lung lobes and nasal cavity tissue were harvested and the tissues were homogenized in DPBS using a mini-beadbeater with 0.1 mm zirconia beads (Biospec catalog no. 11079110zx). The number of CFU was determined by plating dilutions of tissue homogenates on BG blood agar and enumerating the number of colonies per tissue after 72 hours growth at 37 °C. All animal studies were performed in accordance with the recommendations in the Guide for the Care and Use of Laboratory Animals of the NIH. Our protocols were approved by the University of North Carolina IACUC (22-140). All animals were anesthetized for inoculations, monitored regularly, and euthanized properly and humanely. All efforts were made to minimize suffering.

## 8 ACKNOWLEDGEMENTS

We thank Dr. Zachary DeMars for his helpful insight and assistance in preliminary protein purification and biochemical experiments and Luke Vass for his insight and help in investigating PlrR reactions with small molecule phosphodonors. With the help and mentorship of Dr. Nathan Nicely, the mass photometry work was performed in the UNC Protein Expression and Purification & Macromolecular Crystallography core laboratory supported by the National Cancer Institute of the National Institutes of Health under award number P30CA016086.

This work was funded by National Institutes of Health grants GM050860 to RBB and AI129541 and AI153160 to PAC. The content is solely the responsibility of the authors and does not necessarily represent the official views of the National Institute of General Medical Sciences, the National Institute of Allergy and Infectious Disease, or the National Institutes of Health.

## 9 Author Contributions

i. **the conception or design of the study:**
  SAB, ENK, PAC, RBB
ii. **the acquisition, analysis, or interpretation of the data:**
  SAB, ENK, RMJ, LSM, RJO, PAC, RBB
iii. **writing of the manuscript:**
  SAB, ENK, PAC, RBB

## 11 ABBREVIATED SUMMARY

*Bordetella bronchiseptica* encodes a putative two-component system: a sensor histidine kinase termed PlrS and a response regulator termed PlrR. PlrS and PlrR are critical for virulence and persistence within the lower respiratory tract of a murine host. We showed that PlrS exhibited both autokinase activity and phosphatase activity towards PlrR, that PlrS could serve as a phosphodonor for PlrR, and that PlrS phosphatase activity was critical to persistence in the lower respiratory tract.

