## Supplementary material for "Phosphorylation chemistry of the *Bordetella* PlrSR TCS and its contribution to bacterial persistence in the lower respiratory tract": Figures S1 and S2, Tables S1-S3

Submitted to *Molecular Microbiology* 10-06-22

1. **Figure S1.** Autophosphorylation of longer N-terminal ΔHAMPΔPAS proteins
2. **Figure S2.** Phosphatase activity of longer N-terminal ΔHAMPΔPAS proteins
3. **Table S1.** Strains used in this study
4. **Table S2.** Plasmids
5. **Table S3.** Primers

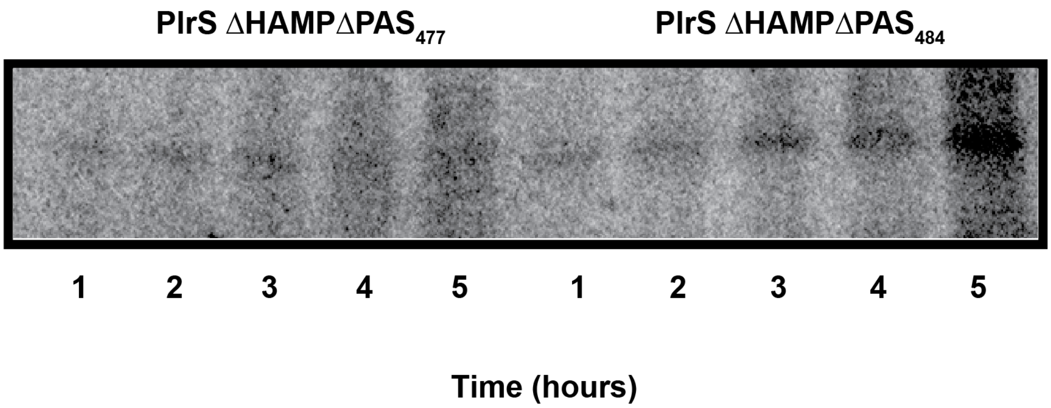

**Figure S1. Autophosphorylation of longer N-terminal ΔHAMPΔPAS proteins.** Phosphorimage of autophosphorylation experiments for PlrS $\Delta$HAMP$\Delta$PAS$_{477}$ and PlrS $\Delta$HAMP$\Delta$PAS$_{484}$ proteins at room temperature over 5 hours.

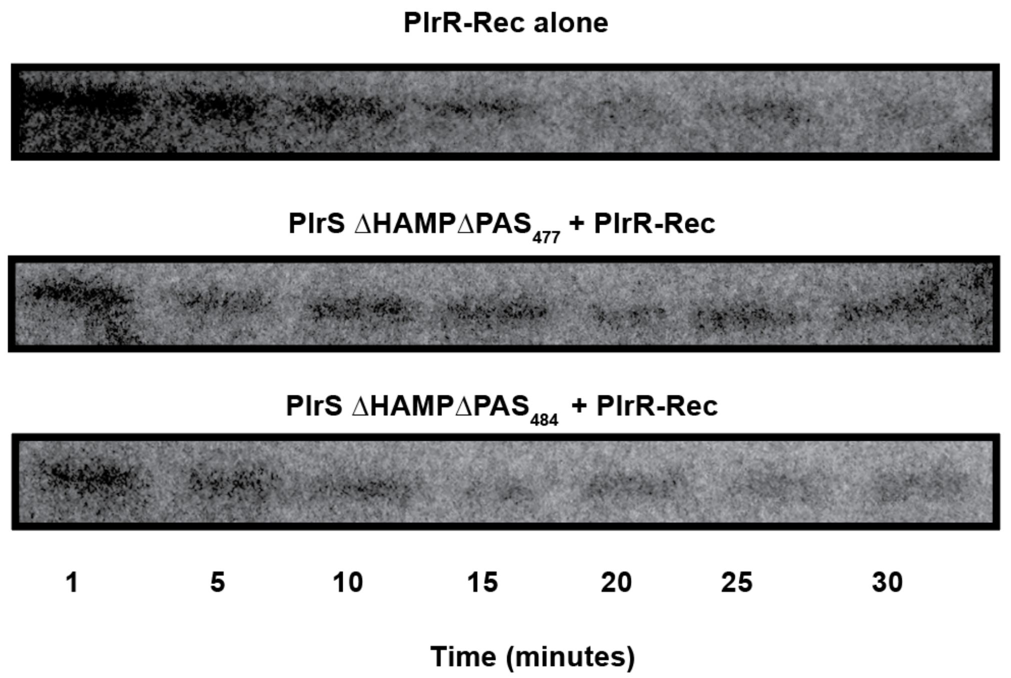

**Figure S2. Phosphatase activity of longer N-terminal ΔHAMPΔPAS proteins.**

Phosphorimages of phosphatase experiments for PlrS ΔHAMPΔPAS$_{477}$ and PlrS ΔHAMPΔPAS$_{484}$ constructs towards PlrR-Rec at room temperature over 30 minutes.

**Table S1.** Strains used in this study

| **Bacterial strain** | **Description** | **Source** |
| --- | --- | --- |
| ***E. coli* strains** | | |
| NEB5α | Molecular cloning strain | NEB |
| DH5α | Molecular cloning strain | Gibco BRL |
| RHO3 | Conjugation strain; *Δasd ΔaphA*, DAP auxotroph | (Lopez *et al.*, 2009) |
| BL21 Star | $F^-$ *ompT* *hsdS*$_B$ ($r_B^-$,$m_B^-$) *gal dcm rne*131 λ(DE3) | Invitrogen |
| BL21 Star/pET28a (+) PlrS$_{316}$ | $Km^r$ | This study |
| BL21 Star/pET28a (+) PlrS$_{316}$ H521Q | $Km^r$ | This study |
| BL21 Star/pET28a (+) PlrS$_{316}$ E522A | $Km^r$ | This study |
| BL21 Star/pET28a (+) PlrS$_{316}$ N525A | $Km^r$ | This study |
| BL21 Star/pET28a (+) PlrS ΔHAMP$_{359}$ | $Km^r$ | This study |
| BL21 Star/pET28a (+) PlrS ΔHAMP$_{359}$ N525A | $Km^r$ | This study |
| BL21 Star/pET28a (+) PlrS ΔHAMPΔPAS$_{494}$ | $Km^r$ | This study |
| BL21 Star/pET28a (+) PlrS ΔHAMPΔPAS$_{494}$ N525A | $Km^r$ | This study |

| | | |
|---|---|---|
| BL21 Star/pET28a (+) PlrS $\Delta$HAMP$\Delta$PAS$_{477}$ | Km$^r$ | This study |
| BL21 Star/pET28a (+) PlrS $\Delta$HAMP$\Delta$PAS$_{484}$ | Km$^r$ | This study |
| BL21 Star/pET28a (+) PlrR | Km$^r$ | This study |
| BL21 Star/pET28a (+) PlrR-Rec | Km$^r$ | This study |
| BL21 Star/pET28a (+) PlrR D52A | Km$^r$ | This study |
| ***B. bronchiseptica* strains** | | |
| RB50 | Wild-type *B. bronchiseptica* strain; Sm$^r$ | (Cotter & Miller, 1994) |
| Δ*plrR att*Tn*7*::*plrR* | RB50 containing a copy of *plrR* at the *att::*Tn7 site driven by the S12 promoter and an in-frame deletion in *plrR* at the native locus; Sm$^r$ Km$^r$ | This study |
| Δ*plrS*$_{5-198}$ Δ*plrR att*Tn*7*::*plrR* | RB50 containing copy of *plrR* at the *att::*Tn7 site driven by the S12 promoter, in-frame deletion in *plrR* at the native locus, and an in-frame | This study |

| | | |
|---|---|---|
| | deletion in *plrS* removing the codons for amino acids 5–198, Sm$^r$ Km$^r$ | |
| *plrS N525A* ∆*plrR att*Tn*7*::*plrR* | RB50 containing a copy of *plrR* at the *att::*Tn7 site driven by the S12 promoter, an in-frame deletion in *plrR* at the native locus and a N525A point mutation in *plrS*; Sm$^r$ Km$^r$ | This study |

**Table S2.** Plasmids

| **Plasmids** | **Description** | **Source** |
|---|---|---|
| pET28a (+) | Protein expression plasmid for *E. coli* expression strains; Encodes an inducible T7 promoter and a N-terminal 6X-His Tag; Km$^r$ | Twist Bioscience |
| pMD11 | pSS4245-based allelic exchange vector for deleting the codons for amino acids 5–198 of *plrS*; Km$^r$ Ap$^r$ | (Kaut *et al.*, 2011) |
| pEG7s | Suicide plasmid for *B. bronchiseptica* used to generate in-frame deletions; Gm$^r$ Ap$^r$ | (Cotter & Miller, 1994) |
| pTNS3 | Tn7 transposase expression vector containing *tnsABCD*; Ap$^r$ | (Anderson *et al.*, 2012) |
| pUCS12 | pUC18-mini-Tn7-km used to integrate genes of interest driven by the S12 promoter to the *att::*Tn7 site; Km$^r$ Ap$^r$ | (Anderson *et al.*, 2012) |
| pMAB09 | pEG7s-based allelic exchange vector for deleting *plrR*; Gm$^r$ Ap$^r$ | This study |
| *plrS N525A*::pEG7s | pEG7s-based allelic exchange vector for introducing a N525A point mutation in *plrS*; Gm$^r$ Ap$^r$ | This study |
| pUCS12-*plrR* | pUCS12-based complementation plasmid that integrates at the *att::*Tn7 site and contains the *plrR* ORF driven by the constitutively active S12 promoter; Km$^r$ Ap$^r$ | This study |

**Table S3.** Primers

| Primer | Sequence |
|---|---|
| PlrS H521Q Fwd | CGTCTGGCGCAGGAAATCAAGAACCCGC |
| PlrS H521Q Rev | GTTCTTGATTTCCTGCGCCAGACGACGCGCAAC |
| PlrS E522A Fwd | CACGCGATCAAGAACCCGCTGACCC |
| PlrS E522A Rev | GATCGCGTGCGCCAGACGACGCGCA |
| PlrS N525A Fwd | GAAATCAAGGCGCCGCTGACCCCGATTCAGC |
| PlrS N525A Rev | GGTCAGCGGCGCCTTGATTTCGTGCGCCAG |
| PlrR D52A Fwd | GACCTGGTGCTGCTGGCTATCTGGATGCCGGAC |
| PlrR D52A Rev | GTCCGGCATCCAGATAGCCAGCAGCACCAGGTC |
